# Non-decision time-informed collapsing threshold diffusion model: A joint modeling framework with identifiable time-dependent parameters

**DOI:** 10.1101/2025.10.30.685574

**Authors:** Amir Hosein Hadian Rasanan, Lukas Schumacher, Michael D. Nunez, Gabriel Weindel, Jörg Rieskamp

## Abstract

Over the past sixty years, evidence accumulation models have emerged as a dominant framework for explaining the neural and behavioral aspects of the process underlying decision making. These models have also been widely used as a measurement instrument to assess individual differences in latent cognitive constructs underlying decision making. A central assumption of most of these models is that decision makers accumulate noisy evidence until a fixed decision threshold is reached. However, both behavioral and neuroscientific findings, along with theoretical considerations related to optimality, have suggested that the decision threshold varies over time. Although time-dependent threshold models often provide a better account of empirical data, a major challenge associated with these models is the unreliable estimation of their parameters. This limitation has led researchers to emphasize model-fitting comparisons rather than interpreting parameter values or accounting for individual differences in the dynamics of the decision threshold. In this work, we address the reliability issue of parameter estimation in time-dependent threshold diffusion models by proposing a joint modeling approach that links non-decision time to external observations. Parameter recovery simulations demonstrate that informing the diffusion model with trial-level noisy measurements of non-decision time substantially improves the reliability of parameter estimation for time-dependent threshold diffusion models. Additionally, we reanalyzed the experimental data from two perceptual decision-making tasks to illustrate the feasibility of the proposed modeling approach. Non-decision time measurements were extracted from electroencephalography (EEG) recordings using the hidden multivariate pattern method. The cognitive modeling results revealed that, in addition to the reliable parameter estimation, constraining non-decision time improves the absolute fit to behavioral data.

## Introduction

For decades, understanding how people make rapid decisions has been a central focus in cognitive science. Studying quick decisions allows researchers to investigate the properties of latent cognitive processes underlying decision making and to develop process-level theories of human choice behavior. Among the various formal cognitive models proposed to explain decision making, evidence accumulation models (EAMs; ***Ratcliff, 1978***; ***Ratcliff and McKoon, 2008***; ***Stone, 1960***; ***Laming, 1968***) represent the most prominent class of cognitive computational models (***Forstmann et al., 2016***). A key advantage of EAMs is their ability to account for both choice and response time (RT) simultaneously (***Forstmann et al., 2016***; ***Ratcliff et al., 2016***). The most widely used variant of EAMs is the well-known diffusion decision model (DDM; ***Ratcliff, 1978***; ***Ratcliff and Rouder, 1998***; ***Ratcliff and McKoon, 2008***), originally proposed to explain behavior in two-alternative choice tasks. The DDM assumes that the decision maker accumulates noisy evidence (with the mean evidence accumulation rate governed by a constant *drift rate*) until the relative evidence for one option reaches a fixed decision threshold. This model has made substantial contributions to our understanding of the neural and cognitive mechanisms underlying decision making (e.g., ***Forstmann et al., 2016***; ***Gold and Shadlen, 2007***; ***Ratcliff et al., 2016***). Moreover, it has been widely employed as a cognitive psychometric instrument for data analysis and for describing latent cognitive processes across various domains, including perceptual decisions (e.g., ***Dutilh and Rieskamp, 2016***; ***Forstmann et al., 2011***; ***Evans and Brown, 2017***), risky decisions (e.g., ***Olschewski et al., 2025***; ***Bhui, 2019***; ***Zhao et al., 2020***), value-based decisions (e.g., ***Krajbich et al., 2010***; ***Fontanesi et al., 2019***; ***Khodadadi et al., 2017***; ***Gluth et al., 2020***), numerical cognition (e.g., ***Ratcliff and McKoon, 2018***; ***Ratcliff, 2022***), intelligence (e.g., ***Schmiedek et al., 2007***; ***Schubert and Frischkorn, 2020***), and aging (e.g., ***Starns and Ratcliff, 2010***; ***Ratcliff, 2008***; ***Ratcliff et al., 2010***; ***von Krause et al., 2022***), among others. Additionally, this model has been employed to investigate the information processing patterns in clinical populations (e.g., ***Nejati et al., 2022***; ***Ging-Jehli et al., 2021***; ***Pirrone et al., 2017***; ***Pedersen et al., 2017***; ***Karalunas et al., 2012***). The broad application of the DDM is attributed to the interpretability of its parameters, which has been validated in several studies (e.g., ***Voss et al., 2004***; ***Arnold et al., 2015***; ***Lerche and Voss, 2019***), as well as their reliability in parameter estimation (***Lerche and Voss, 2017***; ***Yap et al., 2012***).

Although the DDM has become a standard model for describing the latent cognitive constructs of decision making, alternative variants — such as the urgency gating model (***Cisek et al., 2009***; ***Thura et al., 2012***; ***Trueblood et al., 2021***) and collapsing threshold diffusion models (CT-DDMs; ***Drugowitsch et al., 2012***; ***Tajima et al., 2016***) — better align with certain behavioral and neuroscientific findings (***Ratcliff and Frank, 2012***; ***Ging-Jehli et al., 2025***). In particular, the DDM assumes that the decision threshold remains fixed over time, implying that the decision maker requires a constant amount of evidence to make a decision, regardless of the time spent on deliberating. However, several neuroscience studies have shown that human and non-human primates tend to become more urgent as time progresses (e.g., ***Ditterich, 2006***; ***Thura et al., 2012***; ***Cisek et al., 2009***; ***Murphy et al., 2016***; ***Steinemann et al., 2018***; ***Gluth et al., 2012, 2013***; ***Ging-Jehli et al., 2025***; ***Grogan et al., 2025***). These findings support the idea of the collapsing threshold or urgency gating models, in which the required amount of evidence for making a decision decreases over time, thereby increasing the urgency of decision making.^1^

CT-DDMs also predict several behavioral effects that fixed-threshold diffusion models (FT-DDMs) fail to capture. First, CT-DDMs are more flexible in accounting for RT distributions with less asymmetry. Specifically, while FT-DDMs typically predict strongly right-skewed RT distributions (***Ratcliff and Smith, 2004***), CT-DDMs can account for Gaussian-like RT distributions (***Evans and Hawkins, 2019***; ***Hawkins et al., 2015***). Several studies have observed such symmetric, Gaussian-like RT distributions under specific conditions (e.g., ***Roitman and Shadlen, 2002***; ***Ditterich, 2006***; ***Hawkins et al., 2019***; ***Evans and Hawkins, 2019***; ***O’Connell et al., 2012***). A second key prediction of CT-DDMs is that slower responses tend to exhibit greater variability in accuracy (in inferential decisions) or in choice proportion (in preferential decisions) — an effect supported by several empirical studies (e.g., ***Olschewski et al., 2025***; ***Murphy et al., 2016***; ***Steinemann et al., 2018***). In other words, these studies found a higher error rate for trials with longer response times. In contrast, the FT-DDM predicts a constant error rate across both fast and slow responses.^2^ Beyond these behavioral effects, some empirical studies have shown that experimental manipulations such as introducing a decision deadline or emphasizing urgency encourage individuals to adopt a collapsing threshold strategy (***Hawkins et al., 2015***; ***Evans et al., 2020a***; ***Evans and Hawkins, 2019***; ***Katsimpokis et al., 2020***). Furthermore, quantitative model comparison showed broad empirical support for the idea that people adjust their decision threshold during a single decision even without time-pressure manipulation (***Olschewski et al., 2025***; ***Palestro et al., 2018***; ***Bhui, 2019***; ***Khodadadi et al., 2017***; ***Ging-Jehli et al., 2025***; ***Ratcliff and Frank, 2012***).^3^

On top of the supporting evidence for CT-DDMs from empirical studies, there is also a strong theoretical foundation for these models. CT-DDMs offer a normative, optimal policy for evidence accumulation in several scenarios. The FT-DDM maximizes accuracy for a fixed mean RT — or, equivalently, minimizes mean RT for a fixed accuracy level — in environments with homogeneous choice difficulty and no external biases (***Bogacz et al., 2006***; ***Moran, 2015***). However, this information processing strategy is not optimal in more complicated settings. For instance, in environments where the difficulty of the choice problem is unknown or varies across trials (***Drugowitsch et al., 2012***; ***Tajima et al., 2016***; ***Fudenberg et al., 2018***), or where a stochastic deadline governs decision timing (***Frazier and Yu, 2007***), or where external biases (e.g., prior information about the correct response) are present (***Moran, 2015***), CT-DDMs provide the optimal evidence accumulation policy.

Although there is both theoretical and empirical support for CT-DDMs, a major limitation that restricts their application as measurement models for exploring individual differences is that some CT-DDM parameters cannot be reliably estimated. Several systematic evaluations using various parameter estimation methods have reported poor parameter recovery for CT-DDMs (***Evans et al., 2020b***; ***Murrow and Holmes, 2024a***; ***Fengler et al., 2021***), especially when the threshold declines nonlinearly. In other words, these studies found that the actual generating parameters of CT-DDMs are not recoverable. Yet, reliable parameter recovery is essential for interpreting the parameters of a cognitive model. Accordingly, the poor parameter recovery of CT-DDMs prevents the use of parameter estimates to test hypotheses about latent cognitive constructs related to threshold dynamics in decision making (***Kruschke and Liddell, 2018***). Thus, developing a reliable method for estimating CT-DDM parameters is a crucial step toward understanding how individual differences influence within-trial threshold dynamics. This issue is particularly critical given that the conditions under which individuals adjust their decision thresholds during a single trial are still not well understood.

Since CT-DDMs lack a closed-form likelihood function, most methodological efforts to improve parameter estimation have focused on developing numerical procedures to approximate the likelihood function. One such approach is the integral equation method (e.g., ***Buonocore et al., 1987, 1990***; ***Smith, 2000***; ***Zhang et al., 2014***; ***Smith and Ratcliff, 2022***; ***Hadian Rasanan et al., 2025***), which is both simple and computationally efficient. In this method, the first-passage time distribution of the diffusion process is approximated by solving a linear Volterra integral equation of the second kind. Another prominent approach involves partial differential equations, where the first-passage time distribution is estimated by numerically solving a forward or backward Kolmogorov equation (e.g., ***Hadian Rasanan et al., 2023, 2024b***; ***Murrow and Holmes, 2024b***; ***Shinn et al., 2020***; ***Boehm et al., 2021***; ***Richter et al., 2023***; ***Voss and Voss, 2008***). In addition to these two numerical methods, which directly estimate the first-passage time distribution, there are simulation-based techniques that approximate the likelihood function using large-scale simulations. These include kernel density estimation approaches (e.g., ***Turner and Sederberg, 2014***; ***Holmes, 2015***) and neural network-based methods (e.g., ***Fengler et al., 2021***; ***Radev et al., 2023a, 2020***). Each of these likelihood approximation procedures has its strengths and limitations. However, it is important to note that none of these methods fully resolved the reliability issues associated with parameter estimation in CT-DDMs. One reason that more precise likelihood approximation methods did not improve the reliability in parameter estimation is the trade-off between non-decision time and threshold parameters. In other words, there is an identifiability issue in CT-DDMs, which makes estimating threshold dynamics challenging and cannot be resolved by a more precise likelihood approximation. This issue will be discussed further in the next section.

A less explored yet promising solution for improving the reliability of parameter estimation in CT-DDMs is neural-informed cognitive modeling (also known as joint models). Traditionally, these models have been employed to simultaneously fit and predict both behavioral and neural data (e.g., ***Forstmann and Wagenmakers, 2015***; ***Schall, 2004***; ***Turner et al., 2017, 2015***; ***Nunez et al., 2017***; ***Ghaderi-Kangavari et al., 2022, 2023***). Recent studies have demonstrated that neural-informed approaches can also enhance parameter recovery in EAMs and even render previously unidentifiable parameters identifiable (***Nunez et al., 2025***; ***Ghaderi-Kangavari et al., 2023***). Fundamentally, linking DDM parameters to external sources of information — such as neurophysiological signals — introduces additional constraints on the parameter space, thereby enhancing identifiability (e.g., ***Nunez et al., 2025***; ***Ghaderi-Kangavari et al., 2023***). For instance, ***Nunez et al. (2025***) recently proposed a joint modeling framework to identify the diffusion coefficient (i.e., the noise parameter) in the FT-DDM. By constraining the threshold parameter using external observations (e.g., neural signals), the authors demonstrated that one could simultaneously estimate the drift rate, decision threshold, and diffusion coefficient parameters that cannot be jointly identified using only choice and RT data (***Nunez et al., 2025***).

In this work, we explain how non-decision time and collapsing threshold parameters trade off and introduce a non-decision time–informed (NDT-informed) diffusion modeling framework that links non-decision time to external observations, enabling more reliable estimation of collapsing threshold parameters. In contrast to previous methodological studies that primarily focused on likelihood approximation techniques, our contribution lies in proposing a novel *joint modeling* framework, rather than a new estimation algorithm. We demonstrate that informing the model with trial-level noisy measurements of non-decision time — potentially derived from neural data (e.g., ***Nunez et al., 2019***) — substantially improves parameter recovery in CT-DDMs. In addition, linking non-decision time to external observations enhances the model’s fit to behavioral data (i.e., RTs and choices).

The remainder of this paper is organized as follows. First, we present the NDT-informed diffusion modeling framework and discuss a method for estimating non-decision time at the trial level using neural signals, along with theoretical justifications for how constraining non-decision time can improve parameter estimation in CT-DDMs. Next, we report results from an extensive simulation study to assess the effectiveness of the proposed joint modeling framework. Finally, we reanalyze two empirical datasets to illustrate the practical applicability of the method and to examine whether empirical evidence supports the collapsing threshold hypothesis. Finally, we discuss how the proposed NDT-informed diffusion modeling can address the mixed findings in support of CT-DDM and explain how this method can be generalized to other types of diffusion models. We also discuss some behavioral techniques for estimating non-decision time, which do not require neural data.

### Non-decision time-informed diffusion modeling

The evidence accumulation process considered in the diffusion models, including CT-DDM, can be represented by a Wiener process (also known as Brownian motion) fluctuating between two time-dependent thresholds (***Ratcliff et al., 2016***):

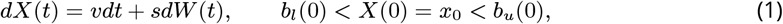

in which *X*(*t*) represents the accumulated evidence until time *t, v* is the mean evidence accumulation rate (drift rate), *s* is the diffusion coefficient, *x*_0_ is the starting point bias (also known as pre-decision bias), and *dW* (*t*) is the Wiener process. The process starts the accumulation from *x*_0_ and continues until the accumulator crosses either the upper (*b*_*u*_(*t*)) or the lower (*b*_*l*_(*t*)) threshold (i.e., *X*(*t*) ≥ *b*_*u*_(*t*) or *X*(*t*) ≤ *b*_*l*_(*t*)). *v* modulates the rate of evidence accumulation, and it corresponds to the quality of the input signal. When the signal-to-noise ratio is high, the task becomes easier, and the speed of evidence accumulation increases. Besides, *s* determines the level of noise in the process. In addition to these components related to decision time, the DDM also includes a non-decision time parameter (*τ* ), which corresponds to the total time unrelated to the decision process, such as perceptual encoding and motor execution. These — namely, *v, x*_0_, *b*_*u*_(*t*), *b*_*l*_(*t*) and *τ* — are the main components of the model. Figure 1 illustrates a fixed threshold and a hyperbolic collapsing threshold.

**Figure 1.**
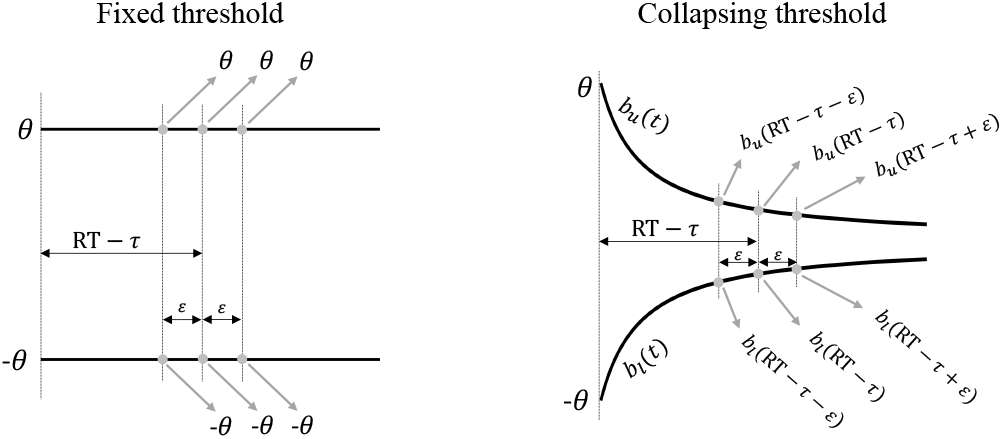
An illustration of the effect of non-decision time on the threshold value on the final stopping point. The illustration of the evidence accumulation process in the fixed threshold diffusion model (left panel; *b*_*u*_(*t*) = −*b*_*l*_(*t*) = *θ*) and hyperbolic collapsing threshold diffusion model (right panel; 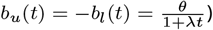.

#### Why constraining non-decision time improves parameter estimation in CT-DDMs

A crucial question that needs to be addressed before presenting the NDT-informed diffusion model is why constraining the non-decision time can enhance the parameter estimation of CT-DDMs. To answer this question, we should note that to compute the likelihood function of the CT-DDMs, we need to incorporate the threshold value in the final stopping time (see the discussion on Girsanov change-of-measure theorem in ***Smith*** (***2016***) and ***Hadian Rasanan et al. (2025***)). The final stopping time determines the decision time or, equivalently, is equal to the difference between the RT and the non-decision time (i.e., *t* = RT − *τ* ). When the decision threshold is fixed over time, its value at the stopping point is the same for any decision time (RT −*τ* ), meaning that non-decision time does not influence the threshold value at the stopping point. However, when the decision threshold is (nonlinearly) time-dependent, the threshold value at the final stopping point depends on non-decision time (i.e., we need to incorporate *b*_*u*_(RT − *τ* ) and *b*_*l*_(RT − *τ* )). In other words, optimizing the likelihood yields maximizing the probability of reaching a particular threshold value at the final stopping point (RT − *τ* ). Therefore, the estimated non-decision time parameter affects the threshold dynamic by forcing the threshold to reach the value at the stopping time that maximizes the likelihood function. Figure 1 illustrates the effect of non-decision time on the threshold value at the final stopping point.

Consequently, the dependence of the threshold value at the final stopping point on non-decision time causes a trade-off between the threshold parameters (e.g., starting threshold and decay rate) and non-decision time. For instance, if the threshold has a hyperbolic dynamic 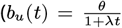, in which *θ* is the starting threshold, and *λ* is the decay rate), the threshold value at the final stopping point becomes equal to 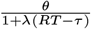 By dividing both the numerator and the denominator by *θ*, we obtain 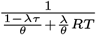, which shows that there is a trade-off between non-decision time, starting threshold, and decay rate. Accordingly, estimating these three parameters simultaneously can be pragmatic. Besides, it implies that inaccurate non-decision time estimation, even if the error is minimal, can lead to erroneous estimates of time-dependent threshold parameters (see Appendix 1 for a simulation study on the tradeoff between non-decision time and starting threshold and decay rate). However, constraining non-decision time in the NDT-informed diffusion modeling framework makes all parameters identifiable, thereby improving the reliability of parameter recovery (see also ***Kira et al., 2025***).

#### Model specification

The traditional diffusion modeling approach estimates model parameters solely based on RT and choice data (e.g., ***Ratcliff and Rouder, 1998***; ***Ratcliff and Tuerlinckx, 2002***; ***Ratcliff and Smith, 2004***). In contrast, in this work, we also assume that an additional source of data related to non-decision time is available. This additional data represents a trial-level noisy measurement of non-decision time, which differs from the actual non-decision time parameter (*τ* ) in the model. Therefore, for each trial *n*, we assume that a measurement of RT_*n*_, Choice_*n*_, and non-decision time measurement (*Z*_*n*_) are available. It is worth highlighting the difference between *Z*_*n*_ and *τ* . *τ* is the mean non-decision time parameter, which is fixed across trials and is estimated through model fitting, whereas *Z*_*n*_ is a trial-level non-decision time measurement (observation) that can be extracted from an external source of data (e.g., from neural signals; see the following section). It is crucial to note that during parameter estimation, we treat *Z*_*n*_ as an input to the model.

Empirical studies on non-decision time measurement usually have reported a right-skewed distribution for their measurements (e.g., ***Weindel, 2021***; ***Weindel et al., 2025***). For instance, the measured perceptual encoding time and motor execution time reported by ***Weindel et al. (2025***) are right-skewed. Therefore, we assume that the observed non-decision time measurements *Z*_*n*_ follow an approximate log-normal distribution, which is right-skewed (we will discuss how this distributional assumption can affect the results later).^4^ Thus, we model the non-decision time measurements *Z*_*n*_, using a log-normal distribution with parameters *µ* and *σ*_*z*_ . The available measurements (i.e., observed data) at trial *n* can be represented as follows:

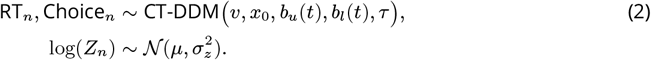

To link the non-decision time measurements (*Z*_*n*_) to the non-decision time parameter (*τ* ), we assume that the actual non-decision time (*τ* ) is the mean value of the observation distribution, which implies 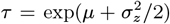 or equivalently 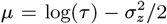. Therefore, the joint negative log-likelihood of the NDT-informed CT-DDM can be formulated as follows:

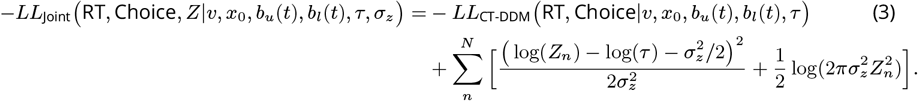

The first term (*LL*_CT-DDM_) corresponds to the likelihood of the choice behavior (i.e., RT and choice data) predicted by the CT-DDM. The second term represents the log-likelihood of the observed non-decision times, which are assumed to follow a log-normal distribution with a mean of *τ* and a shape parameter of *σ*_*z*_ . The non-decision time parameter *τ* is shared across both terms, and linking it to trial-level measurements of non-decision time (*Z*_*n*_) helps constrain its estimation. By minimizing this joint negative log-likelihood function, we can simultaneously estimate the CT-DDM parameters that predict human choice behavior and the summary statistics of the non-decision time observations.

As previously noted, an exact likelihood function for CT-DDMs is not available. Nevertheless, several numerical methods exist for approximating this likelihood, including the integral equation method (e.g., ***Smith, 2000***; ***Zhang et al., 2014***; ***Smith and Ratcliff, 2022***) and the partial differential equation method (e.g., ***Hadian Rasanan et al., 2023***; ***Shinn et al., 2020***; ***Boehm et al., 2021***; ***Richter et al., 2023***). Any of these methods can be used to compute −*LL*_CT-DDM_. In this work, we employed the integral equation method, which is a relatively simple and accurate approach (***Richter et al., 2023***; ***Hadian Rasanan et al., 2025***). In Appendix 2, we detail the procedure for computing −*LL*_CT-DDM_ using the integral equation method. Moreover, we replicated the main results using a BayesFlow approach (***Radev et al., 2020***) to assess sensitivity to the likelihood approximation method.

#### How to measure non-decision time

The central assumption of the NDT-informed diffusion modeling is that the non-decision time measurement (*Z*_*n*_) is available for each trial. Thus, providing a reliable measurement of non-decision time is crucial for the proposed model. Measuring non-decision time using neurophysiological signals like electromyographic (EMG) or electroencephalogram (EEG) has been studied in the field of model-based cognitive neuroscience for a decade (e.g., ***Servant et al., 2016***; ***Nunez et al., 2019***; ***Weindel et al., 2021a***; ***Ghaderi-Kangavari et al., 2022***). Accordingly, several techniques have been proposed in the literature to measure non-decision time. One class of methods involves measuring single-trial event-related potentials (ERPs), enabling researchers to link the ERP components to cognitive parameters of the diffusion model through a (non-)linear regressor (e.g., ***Nunez et al., 2017***; ***Bridwell et al., 2018***; ***Ghaderi-Kangavari et al., 2022, 2023***). One main single-trial ERP component of interest is the N200 peak latency, which is thought to correlate with perceptual encoding time (***Nunez et al., 2019***). However, single-trial N200 latencies capture only a portion of non-decision time, which relates to perceptual encoding, and thus, cannot be used directly as a measurement of total non-decision time. Furthermore, incorporating the N200 component as a regressor for the non-decision time parameter does not resolve the identifiability issue of the CT-DDM. This limitation was highlighted by ***Ghaderi-Kangavari et al. (2023***), who conducted a systematic study on parameter recovery for various neural-informed diffusion models. They reported unreliable parameter estimates for the CT-DDM in which non-decision time was linked to trial-level N200 latency via a regressor (see models 5 and 6 in ***Ghaderi-Kangavari et al., 2023***). The poor parameter recovery in their models arises from a trade-off between the threshold parameters and the regression coefficients — a problem similar to the identifiability issue we discussed earlier. Therefore, informing the model with N200 latency does not resolve the underlying identifiability problem.

Similarly, an EMG signal can be used to measure the execution time of motor responses (***Weindel et al., 2021a***). In this approach, we can detect and measure the motor execution time based on the muscle activity. Although this method provides an accurate measure of motor execution, it does not capture total non-decision time and fails to address the parameter recovery issues in CT-DDMs reported by ***Ghaderi-Kangavari et al. (2023***). Furthermore, recent evidence shows that at least some portion of the EMG signal is related to evidence accumulation (***Servant et al., 2021***; ***Weindel, 2021***; ***Weindel et al., 2021b***). Although a potential approach would be to use both EMG and EEG signals together to estimate perceptual encoding and motor execution time, this approach is relatively complicated.^5^

More recent advancements in neural signal analysis have led to the development of methods to discover single-trial EEG or MEG events across the entire trial-level time series (e.g., ***Anderson et al., 2016***; ***Weindel et al., 2024***). These methods provide the sequence and timing of latent cognitive steps at each trial and have already been used to uncover the cognitive stages in various empirical paradigms, including perceptual decision tasks (***Van Maanen et al., 2021***), lexical decision tasks (***Berberyan et al., 2021***; ***Krause et al., 2024***), and associative recognition tasks (***Van Maanen et al., 2021***). In this paper, we have employed the hidden multivariate pattern (HMP) method (***Weindel et al., 2024***) to extract non-decision time from the EEG signal. It has been demonstrated that this method can decompose the RT into components associated with decision-relevant features versus decision-unrelated physical properties of the stimuli (***Weindel et al., 2025***). Thus, we can extract the decision time from neural signals, and by subtracting the estimated decision time from observed RT, we can obtain a measurement of non-decision time in each trial. The entire procedure of NDT-informed diffusion modeling, which involves extracting non-decision time from neural signals using the HMP method, is presented in Figure 2.

**Figure 2.**
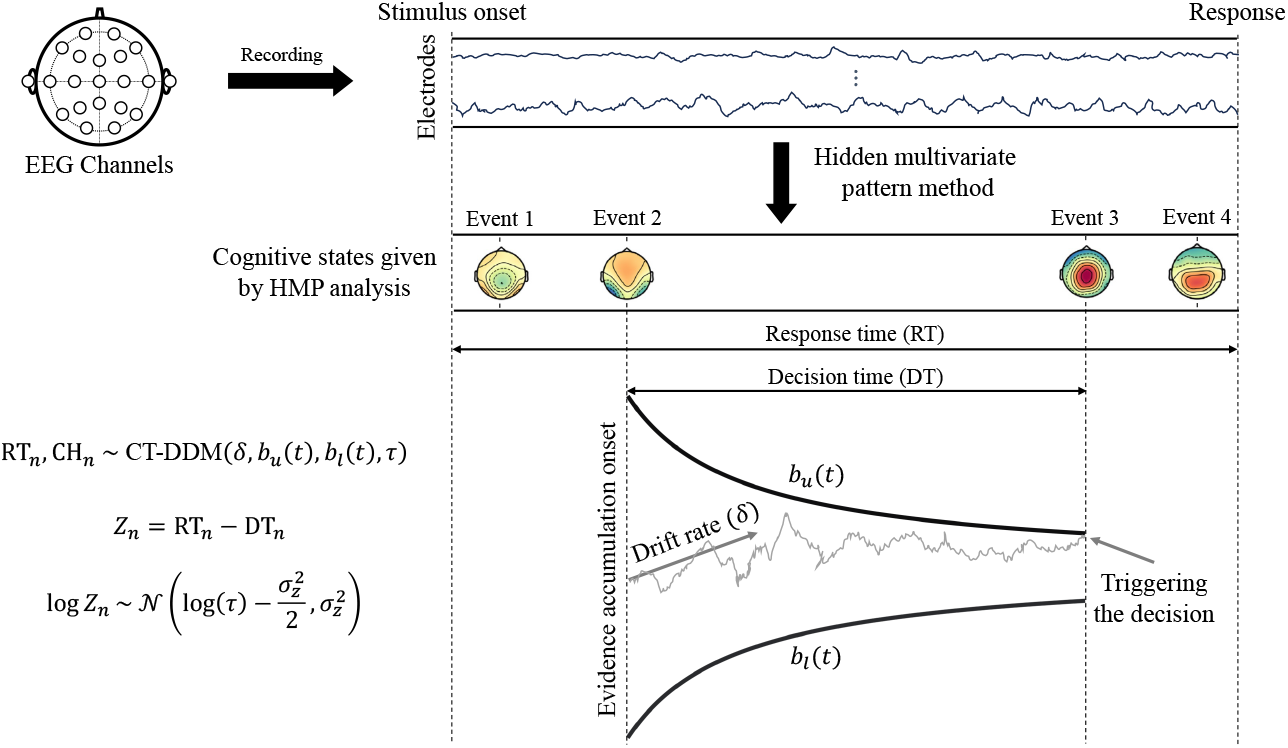
Illustration of the non-decision time-informed diffusion model. The HMP analysis method extracts the timing of cognitive states in each trial from EEG signals. Then, the decision time is considered the duration between the cognitive states that determines the decision process, such as the N200 latency, which marks the end of perceptual encoding (***Nunez et al., 2019***), and the peak of central parietal positivity, which indicates the end of the decision process (***O’Connell et al., 2012***) — see also ***Weindel et al. (2025***) for a detailed discussion. Subtracting the decision time from the response time gives an approximation of non-decision time in each trial. The extracted non-decision time measurements are employed to constrain the non-decision time parameter in the CT-DDM.

### Simulation Study 1: Parameter recovery

#### Methods

We conducted a parameter recovery study using simulated data to evaluate whether the parameters of the NDT-informed CT-DDM can be estimated reliably. To assess the robustness of our method, we considered two distinct nonlinear functional forms for the threshold dynamics. Theoretical studies on optimality have shown that the optimal decision threshold follows a nonlinear, monotonically decreasing trajectory (***Fudenberg et al., 2018***; ***Frazier and Yu, 2007***). These studies suggest that hyperbolic or exponential functions are the most plausible candidates for modeling the optimal threshold (***Fudenberg et al., 2018***; ***Frazier and Yu, 2007***). These collapsing thresholds have already been used in several studies (e.g., ***Olschewski et al., 2025***; ***Bhui, 2019***; ***Milosavljevic et al., 2010***; ***Voskuilen et al., 2016***; ***Hanks et al., 2011***). Additionally, ***Smith and Ratcliff*** (***2022***) demonstrated that an urgency gating model with a linear urgency signal is mathematically equivalent to a hyperbolic CT-DDM (see also ***Trueblood et al., 2021***). It is also important to note that prior research has reported poor parameter recovery for nonlinear collapsing threshold models (***Evans et al., 2020b***; ***Murrow and Holmes, 2024a***). Therefore, we selected the exponential collapsing threshold diffusion model (ECT-DDM) and the hyperbolic collapsing threshold diffusion model (HCT-DDM) — two of the most theoretically motivated and widely supported nonlinear threshold dynamics — for our simulations:^6^

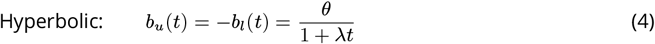

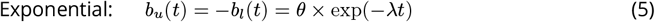

where *θ* is the starting threshold (i.e., *b*_*u*_(0) = *θ*) and *λ >* 0 is the decay rate. *b*_*u*_(*t*) stands for the upper threshold, and the lower threshold is equal to the reflection of the upper threshold (i.e., *b*_*l*_(*t*) = −*b*_*u*_(*t*)). In addition to these two non-linear collapsing threshold dynamics, we also considered linear collapsing threshold models, defined as *b*_*u*_(*t*) = −*b*_*l*_(*t*) = *θ* − *λ t*, to check the robustness of the results. The results were largely consistent with the non-linear models, as detailed in Appendix 3. In all simulations, we assumed that the accumulation process starts from zero (i.e., starting point *x*_0_ = 0), which implies that there is no *a priori* bias towards one option in the simulation data. Also, we set the diffusion coefficient equal to one (i.e., *s* = 1). To simulate RT and choice, we considered the following discrete form of the accumulation process (1) with a time step Δ*t* = 0.001:

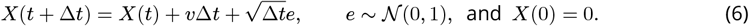

To cover all possible model behavior in the simulations, we used the following uniform distributions for the CT-DDM parameters (i.e., *δ, τ, θ*, and *λ*) and the shape parameter of the non-decision time measurements (*σ*_*z*_ ):^7^

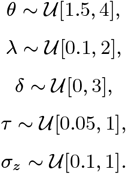

After sampling random parameters from the uniform distributions described above, we generated RT, choice, and random non-decision time measurements (*Z*_*n*_) for various trial numbers (i.e., 100, 250, 500, 750, and 1000). For each number of trials, we generated 1000 datasets. As we already mentioned, we assumed that the actual non-decision time (*τ* ) is assumed to be equal to the mean value of the log-normally distributed non-decision time measurements. Thus, the location parameter of the log-normal distribution is obtained by 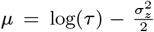. Therefore, in each trial, we sampled a noisy observation of non-decision time using the following log-normal distribution:

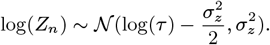

We have employed the differential evolution optimization routine to minimize the joint negative log-likelihood (3). Also, to approximate the likelihood function of the CT-DDM, we used the integral equation method with a time step Δ*t* = 0.02 (see Appendix 2 for the details of likelihood approximation). A problem with the maximum likelihood procedure is its sensitivity to outliers, which can yield likelihood values of zero (***Ratcliff and Tuerlinckx, 2002***). Outliers can affect the fitting procedure and cause convergence problems (***Ratcliff and Tuerlinckx, 2002***; ***Heathcote et al., 2002***). We truncated the likelihood values lower than 10^−14^ to address this issue. To account for potential influences of the estimation method on parameter inference, we replicated the main results using amortized Bayesian inference as implemented in BayesFlow (***Radev et al., 2020, 2023b***). The obtained results based on this alternative approach are presented in Appendix 4.

To evaluate the precision of parameter recovery, we employed the R-squared (*R*^2^) index, which is defined as follows:

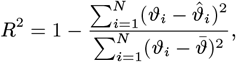

in which *ϑ* and 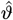 stand for the actual generating parameter and estimated parameter, respectively, and 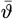 in the denominator shows the mean of the true parameter. Based on the *R*^2^ index, we can assess how well the estimated parameters align with the actual data-generating parameters. This metric has already been employed in the literature to evaluate parameter estimation methods and assessment of estimation reliability in cognitive models (e.g., ***Radev et al., 2020***; ***Hadian Rasanan et al., 2025***).

#### Results

Figure 3 illustrates the *R*^2^ value for the threshold parameters (i.e., starting threshold *θ*, and decay rate *λ*) for both the NDT-informed and uninformed CT-DDM, based on 500 trials. First, this plot demonstrates a substantial improvement in the reliability of threshold parameter recovery in the NDT-informed CT-DDM compared to the uninformed CT-DDM. This improvement is more pronounced for the hyperbolic threshold than the exponential threshold. Second, the results indicate that threshold parameter estimation is generally reliable for both the hyperbolic and exponential models, with particularly high precision observed in the exponential case.

**Figure 3.**
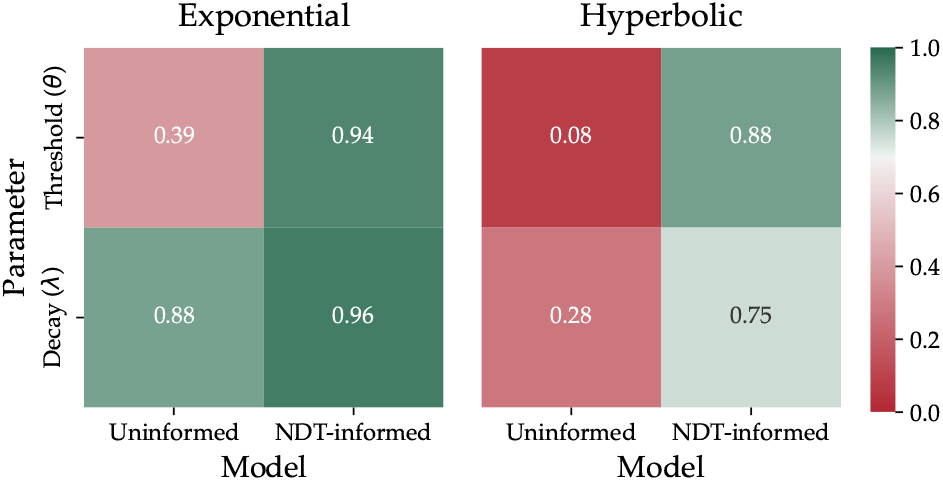
*R*^2^ values measuring the agreement between estimated and ground truth threshold parameters in exponential (left) and hyperbolic (right) collapsing threshold models.

Figure 4 shows the estimated parameters against the true parameters for both NDT-informed CT-DDMs and uninformed CT-DDMs based on 500 trials. The results suggest that the starting threshold parameter in both hyperbolic and exponential uninformed CT-DDM is unrecoverable, which aligns with previous parameter recovery assessment studies (***Evans et al., 2020b***; ***Murrow and Holmes, 2024a***; ***Ghaderi-Kangavari et al., 2023***). Additionally, the decay rate in the hyperbolic uninformed CT-DDM exhibits poor parameter recovery. In contrast, the parameters estimated by the NDT-informed CT-DDM closely match the true data-generating parameters. The only challenging parameter to estimate is the decay rate in the hyperbolic CT-DDM. As shown in Figures 3 and 4, recovering this parameter is more challenging in the hyperbolic model compared to the exponential CT-DDM.

**Figure 4.**
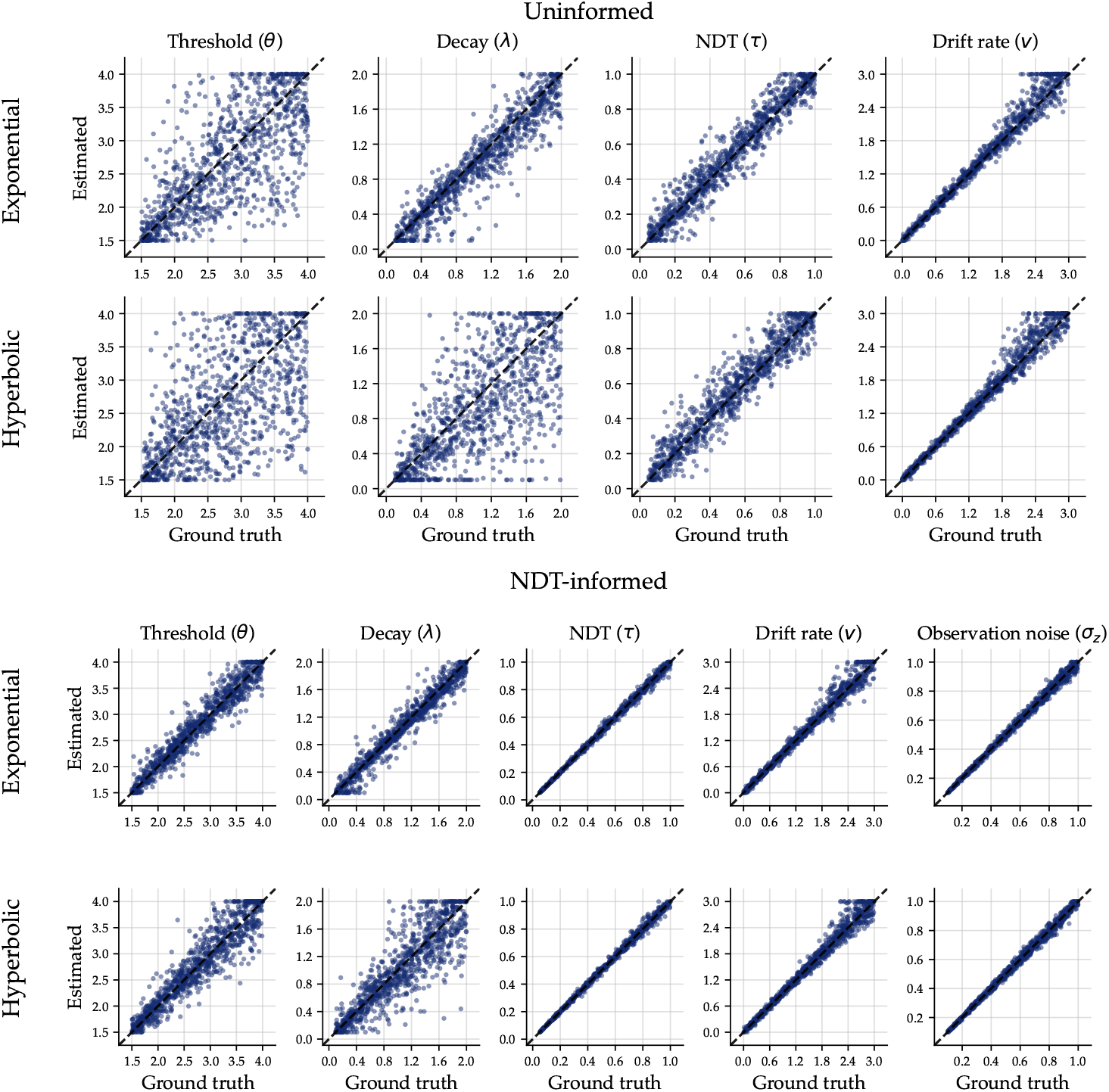
Estimated versus ground truth parameter values for two modeling approaches. The top two rows show parameter recovery for models estimated without additional constraints (“Uninformed”), while the bottom two rows show recovery for joint models that incorporate additional non-decision time observations (“NDT-informed”). Each point represents a recovered parameter estimate plotted against its corresponding true data-generating value. Rows correspond to different functional forms of threshold (i.e., Exponential and Hyperbolic). Dashed diagonal lines indicate perfect recovery.

Figure 5 illustrates the sensitivity of parameter estimation in both NDT-informed and uninformed CT-DDMs as a function of the number of trials. As expected, increasing the number of trials generally improves parameter recovery. However, in the uninformed approach, the collapsing threshold parameters cannot be reliably estimated even with 1000 trials. In contrast, NDT-informed CT-DDMs achieve reliable parameter recovery with substantially fewer trials. Specifically, in the NDT-informed hyperbolic model, accurate estimation of the decay rate requires more than 250 trials. In the exponential NDT-informed model, both the starting threshold and decay rate can be reliably estimated with as few as 100 trials. The drift rate and non-decision time parameters are estimated with high precision, even with 100 trials in both informed and uninformed models, regardless of whether the threshold follows a hyperbolic or exponential form.

**Figure 5.**
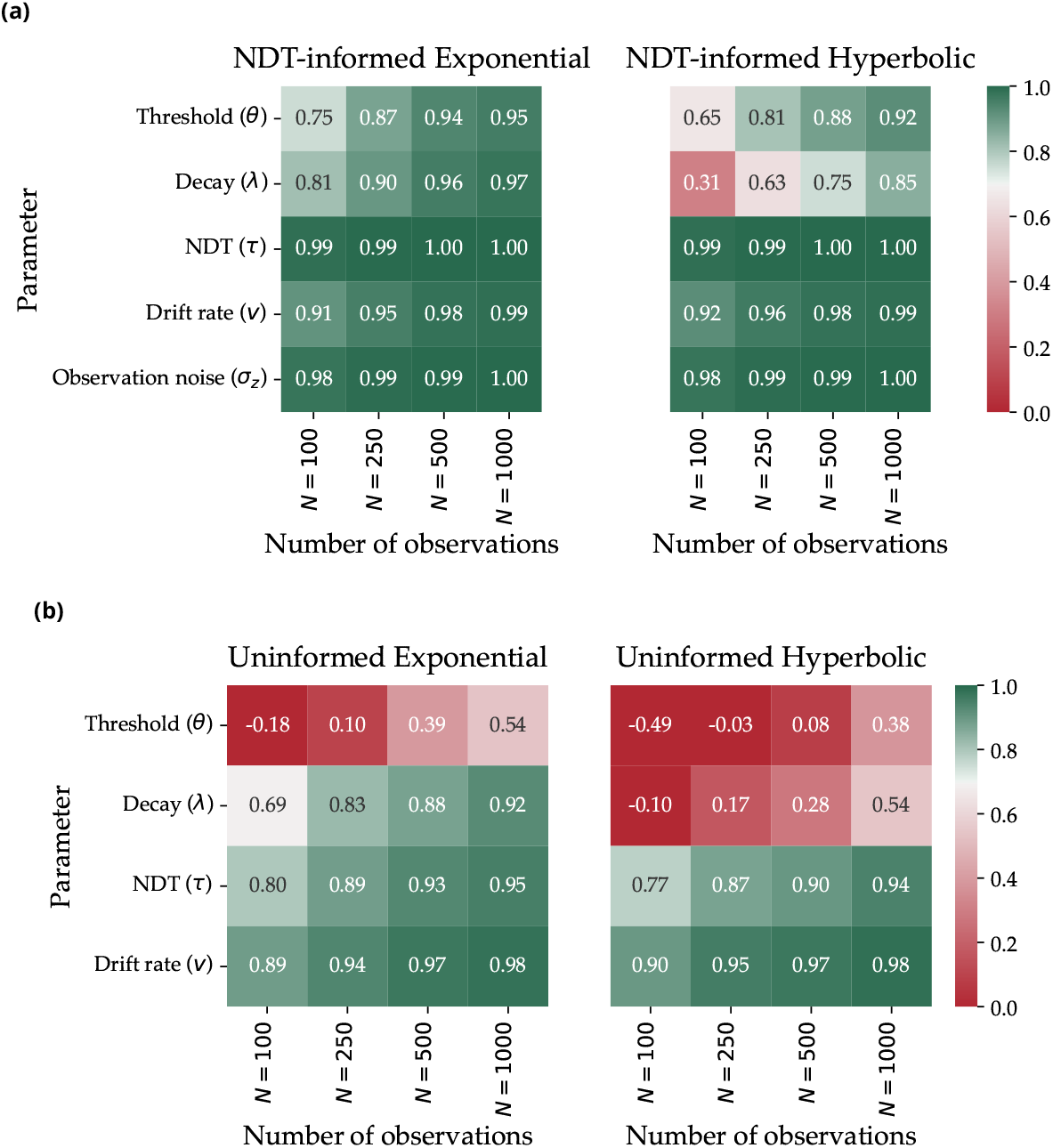
Illustration of the sensitivity of parameter estimation to the number of trials in the NDT-informed (a) and uninformed models (b). *R*^2^ values measure the agreement between estimated and ground truth parameters of the exponential (left) and hyperbolic (right) collapsing threshold models.

Figure 6 presents the sensitivity of the parameter estimation in NDT-informed CT-DDM to the noise level in non-decision time observation (*Z*_*n*_). We simulated data using three fixed levels of standard deviation for the non-decision time measurements (i.e., SD = 0.3, 0.6, and 0.9 seconds), with 500 trials generated for each noise level. The results demonstrate that even with a high level of variability (noise) in non-decision time measurements, parameter estimation remains reliable. Moreover, these results demonstrate that the parameter recovery of NTD-informed CT-DDMs is better than the parameter estimation of the uninformed CT-DDM, even under high levels of noise in non-decision time measurements.

**Figure 6.**
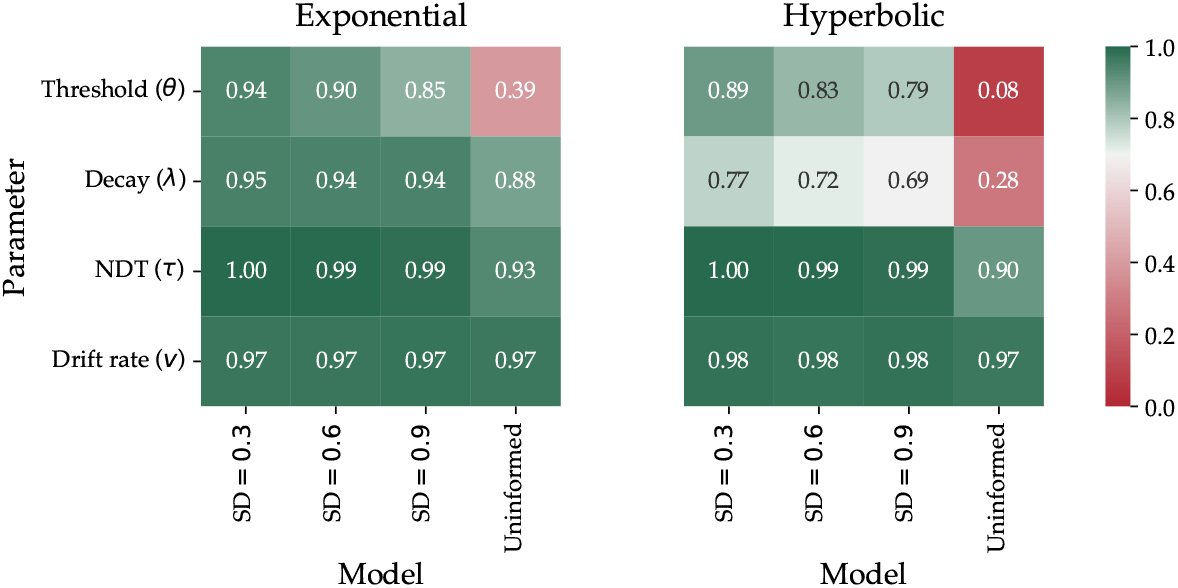
Illustration of the sensitivity of parameter estimation to the noise level in the non-decision time observations. *R*^2^ values measuring the agreement between estimated and ground truth parameters of the exponential (left) and hyperbolic (right) collapsing threshold models.

Additionally, we examined the effect of systematic bias in non-decision time measurement on parameter recovery. Appendix 5 presents the results in the presence of biased non-decision time measurement (i.e., systematically underestimated or overestimated). The results revealed that, even in the presence of biased non-decision time measurement, the actual generating parameters show high correlations with the estimated parameters. Underestimation in non-decision time leads to overestimation in the starting threshold and decay rate. Conversely, overestimation in non-decision time leads to underestimation in both the starting threshold and non-decision time.

### Simulation Study 2: Model recovery

#### Methods

One of the main counterarguments against CT-DDMs is that the slow error responses can also be explained by across-trial variability in drift rate (***Voskuilen et al., 2016***). An FT-DDM that incorporates across-trial variability in drift rate can account for slow errors, and it often achieves fitting performance comparable to that of the CT-DDM when applied to empirical data (***Voskuilen et al., 2016***; ***Smith and Ratcliff, 2022***). This similarity in model performance makes it difficult to determine, based solely on model fit, whether the underlying evidence accumulation process involves a collapsing threshold or a fixed threshold with drift variability. The aim of this section is to show that NDT-informed diffusion modeling enables distinguishing between the CT-DDMs and FT-DDMs with variability in drift rate. To this aim, we conducted a cross-model fitting simulation and a model recovery simulation.

The cross-model fitting simulation is conducted to investigate how parameter estimation in CT-DDMs is affected when the true generative process is an FT-DDM with across-trial variability in the drift rate. In this simulation, we generated data from an FT-DDM with across-trial variability in drift rate and then fitted a CT-DDM without trial-to-trial variability to the simulated data. Across-trial variability in drift rate was modeled as a normal distribution with mean *δ* and standard deviation of *η* (i.e., *v*_*t*_ ∼ 𝒩 (*v, η*), in which *v*_*t*_ is the trial-level drift rate). The following uniform distributions were used to sample the parameters of the FT-DDM:

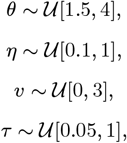

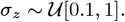

We generated 1000 parameter sets, and for each set, we simulated 500 trials from an FT-DDM with across-trial variability in drift rate. As in the previous simulation study, we assumed that trial-level measurements of non-decision time are available, enabling us to fit NDT-informed diffusion models. Specifically, we fit both an NDT-informed CT-DDM with a hyperbolic collapsing threshold and an NDT-informed CT-DDM with an exponential collapsing threshold to each simulated dataset.

In the model recovery study, we considered two classification problems: a) FT-DDM with drift variability versus exponential CT-DDM and b) FT-DDM with drift variability versus hyperbolic CT-DDM. Specifically, in each classification study, we generated 1000 parameter sets — 500 from an FT-DDM with across-trial variability in drift rate and 500 from the CT-DDM — and simulated 500 trials from each generative model. In each classification study, both the FT-DDM with drift variability and the CT-DDM were then fitted to every simulated dataset. To infer the underlying generating process, we considered two approaches. In one approach, we only considered how well each model was fitted to the simulated data sets, and we compared the goodness-of-fit measures to infer the underlying generating process.^8^ In another approach, we also considered the recovered decay rate (*λ*) value and used this parameter as an additional indicator for the underlying generative process. Therefore, in the second approach, on top of comparing the goodness-of-fits, if the recovered decay rate was lower than 0.1, we classified it as an FT-DDM model.

#### Results

Figure 7 illustrates the parameter recovery results for the model misspecification study.^9^ The most important result is related to the decay rate parameter (*λ*). For more than 75% of the datasets, the estimated decay rate is approximately zero, and only for 5% is the estimated decay rate larger than 0.1. Moreover, these results demonstrate that both the non-decision time parameter and the variability of non-decision time observations can be recovered with high accuracy. The recovered drift rate and starting threshold parameters showed a slight underestimation for larger values. However, the *R*^2^ value remains high for these parameters (i.e., *R*^2^ *>* 0.85).

**Figure 7.**
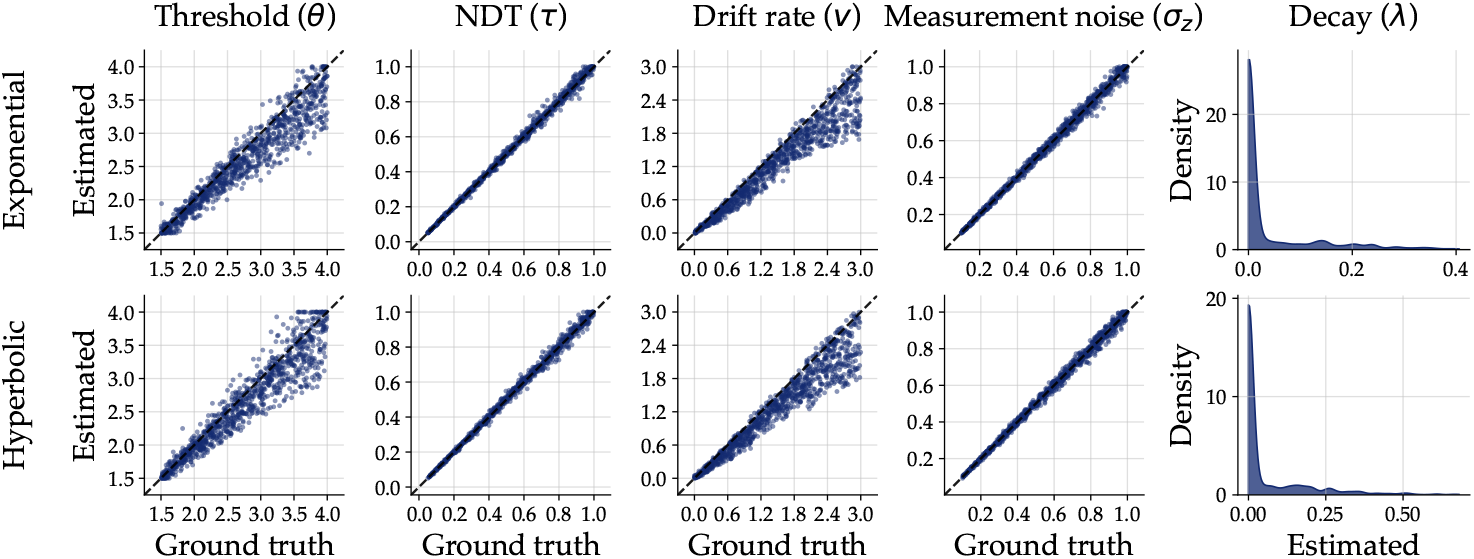
Estimated versus true parameter values for cross-fitting of CT-DDM on FT-DDM with across-trial variability in drift rate. In each panel except the rightmost one, each point represents a recovered parameter estimate plotted against its corresponding true data-generating value, and the dashed diagonal line indicates perfect recovery. The right column illustrates the density of the estimated decay rate parameter (*λ*). Rows correspond to different functional forms of threshold (i.e., Exponential and Hyperbolic).

Table 1 summarizes the model recovery results for both exponential and hyperbolic CT-DDMs. The results indicate that the exponential model exhibits better recovery performance than the hyperbolic model, consistent with its superior parameter recovery. More importantly, incorporating the estimated decay rate into model inference substantially improves model recovery accuracy. In other words, rather than relying solely on goodness-of-fit indices, using the estimated decay rate to guide model classification (i.e., models with an estimated decay rate below 0.1 were classified as FT-DDMs) can improve the precision of model recovery. Because the NDT-informed framework provides reliable estimates of collapsing threshold parameters, these parameter values can be used confidently in model inference, thereby improving the precision of model recovery.^10^

**Table 1.** The model recovery results for exponential and hyperbolic collapsing threshold diffusion models.

|  | Exponential | Hyperbolic |
| --- | --- | --- |
| Goodness of fit only | 89.3% | 86.9% |
| Goodness of fit and decay value | 97.7% | 95.1% |

In sum, the simulation results confirm that incorporating non-decision time information into the diffusion model enables distinguishing between CT-DDM and FT-DDM. Moreover, these results suggest that NDT-informed diffusion modeling can be reliably applied to infer the underlying threshold dynamics. In other words, if the underlying generating process is an FT-DDM (with or without across-trial variability), then the estimated decay rate would be very close to zero or a relatively very small value. Therefore, we can infer the underlying threshold dynamics through the estimated decay rate value in the NDT-informed diffusion modeling framework.

### Applications to empirical data

We present two case studies to: (1) show the feasibility of the proposed NDT-informed diffusion modeling approach, (2) explore whether informing the diffusion model with noisy measurements of non-decision time can enhance model fitting to behavioral data, and (3) investigate whether people adjust the decision threshold within a single trial (at least in the case studies considered here).^11^ To analyze the behavioral data, we consider six different diffusion models. For each dataset, we fitted the NDT-informed HCT-DDM, ECT-DDM, and FT-DDM to compare the fixed threshold against the collapsing threshold using the joint modeling framework. Also, to check for improvements of NDT-informed, we fitted the uninformed version of HCT-DDM and ECT-DDM. Additionally, as a benchmark, we fitted an FT-DDM that includes trial-to-trial variability in drift rate. Then, we employed the Bayesian information criterion (BIC) to evaluate the goodness-of-fit and to compare the models. We compared the considered diffusion models in two ways: with respect to fitting on behavioral data (only RT and choice) and also fitting on all observations, including RT, choice, and non-decision time measurements. Thus, we defined these two BICs for model comparisons:

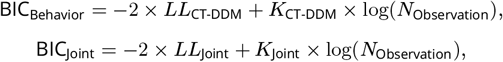

in which *LL*_CT-DDM_ is the log-likelihood of the (collapsing threshold) diffusion model (see Appendix 2 for the details of the likelihood approximation using the integral equation method), *LL*_Joint_ is the joint log-likelihood of behavioral data and non-decision time estimates defined in equation 3, *K*_CT-DDM_ is the number of free parameters in the (collapsing threshold) diffusion model, *K*_Joint_ is the number of free parameters in the joint model, and *N*_Observation_ is the number of observed data points. BIC_Behavior_ only considers the goodness-of-fit on RT and choice, while BIC_Joint_ also considers how well the model can predict the non-decision time observations. Thus, we can compare all six considered models based on BIC_Behavior_ and check whether constraining the non-decision time can improve the fitting of behavioral data. However, BIC_Joint_ is only used for comparing the NDT-informed diffusion models. It is worth noting that comparing joint and behavioral models using BIC_Behavior_ is uncommon and constitutes a limitation of the model comparison study.

#### Study 1: Weindel et al. (2025) dataset

In the first study, we considered data from ***Weindel et al. (2025***). Twenty-six participants took part in the study and performed a binary choice task involving contrast discrimination. On each trial, two Gabor patches were presented, and participants indicated which option had the higher contrast. The difference in contrast level between the two options was always fixed at 5%, while the absolute contrast levels varied from 5% to 95%. In total, participants completed eight experimental blocks, each containing 140 trials. At the beginning of each block, participants received a speed-accuracy instruction. In half of the blocks, participants were requested to focus on decision speed, and in the other half, they were asked to focus on accuracy. The decision difficulty was designed to follow Fechner’s law (***Fechner, 1860***). According to this law, keeping the difference between options constant makes it harder to discriminate between two options as the overall value increases. To easily approximate this law, the following relation between contrast level and drift rate was implemented:

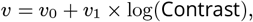

in which *v*_0_, and *v*_1_ are free parameters. *v*_0_ is the baseline drift rate value, and *v*_1_ modulates the sensitivity to the contrast level. The non-decision time measurements were obtained from the fit of an HMP model using the standard cumulative procedure (***Weindel et al., 2024***) on preprocessed EEG data with minimal trial rejection (see ***Weindel et al., 2025***, for the detailed procedure). The HMP model estimated five cognitive events. Their associated single-trial timing and electrode contributions allowed us to interpret these events as relating to early visual processing, perceptual encoding, attention orientation, motor planning, and decision termination. For this analysis, we subtracted the time between the attention orientation (third event) and decision termination (last event) from the RT at each trial to yield a measure of non-decision time. See Figure 8 for the sequence of these events and the time interval between them.

**Figure 8.**
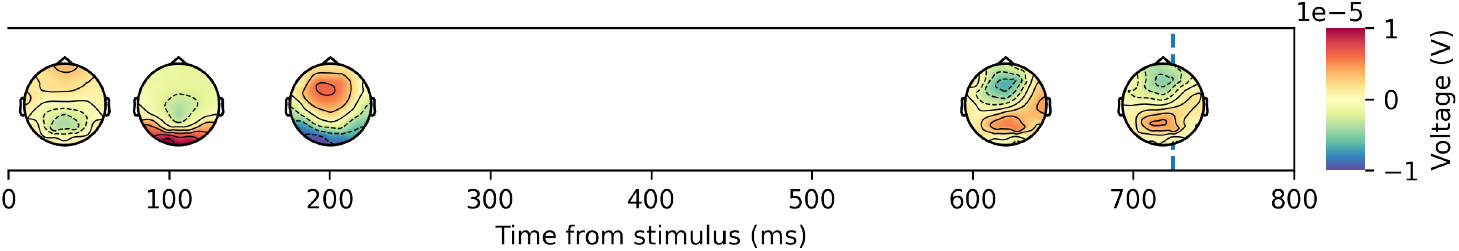
The event sequence and timing of each event estimated by the HMP method averaged over all participants (***Weindel et al., 2025***).

The quantitative results revealed that NDT-informed HCT-DDM outperforms the other models. Table 2 presents the mean estimated parameters of the computational models and a model comparison result based on BIC. The best-fitting model in both speed and accuracy conditions is the NDT-informed HCT-DDM. Also, these results show that both NDT-informed models have better predictive accuracy on RT and choice compared to the two uninformed counterpart models (i.e., based on BIC_Behavior_). The estimated parameters in both NDT-informed models show that the mean starting threshold in the speed condition is lower than in the accuracy condition. However, the mean decay rate is larger in the speed condition compared to the accuracy condition. This implies that the decision threshold starts at a lower point and declines faster in the speed condition than in the accuracy condition.

**Table 2.** The mean estimated parameters and goodness of fit results for Study 1.

| | Model | $v_0$ | $v_1$ | $\tau$ | $\theta$ | $\lambda$ | $\sigma_z$ | BIC<sub>Joint</sub> | BIC<sub>Behavior</sub> |
| --- | --- | --- | --- | --- | --- | --- | --- | --- | --- |
| Speed | NDT-inf HCT-DDM | 2.77 | 0.51 | 0.19 | 1.28 | 2.67 | 0.31 | **-36990.16** | **2644.03** |
|  | NDT-inf ECT-DDM | 2.80 | 0.51 | 0.19 | 1.16 | 1.61 | 0.31 | -36952.86 | 2751.98 |
|  | NDT-inf FT-DDM | 3.01 | 0.54 | 0.21 | 0.69 | — | 0.37 | -29207.05 | 6528.17 |
|  | Uninf HCT-DDM | 2.87 | 0.53 | 0.21 | 1.16 | 2.73 | — | — | 2707.24 |
|  | Uninf ECT-DDM | 2.92 | 0.54 | 0.21 | 1.07 | 1.40 | — | — | 2802.94 |
| Accuracy | NDT-inf HCT-DDM | 3.17 | 0.59 | 0.21 | 1.64 | 1.26 | 0.33 | **-18225.91** | **14919.20** |
|  | NDT-inf ECT-DDM | 3.19 | 0.59 | 0.21 | 1.40 | 0.58 | 0.33 | -17862.53 | 15153.47 |
|  | NDT-inf FT-DDM | 3.11 | 0.57 | 0.24 | 1.00 | — | 0.41 | -9196.26 | 17301.73 |
|  | Uninf HCT-DDM | 3.32 | 0.63 | 0.28 | 1.47 | 1.09 | — | — | 15863.86 |
|  | Uninf ECT-DDM | 3.44 | 0.65 | 0.31 | 1.07 | 0.32 | — | — | 15915.51 |

Note. NDT-inf = NDT-informed and Uninf = Uninformed

Figure 9 demonstrates the estimated threshold dynamics for each individual (gray lines) and the group average (blue lines) by NDT-informed CT-DDMs (see Appendix 6 for the estimated collapsing threshold by uninformed models). The estimated threshold for each individual indicates that most of the participants employed a collapsing threshold strategy (even in the accuracy condition).

**Figure 9.**
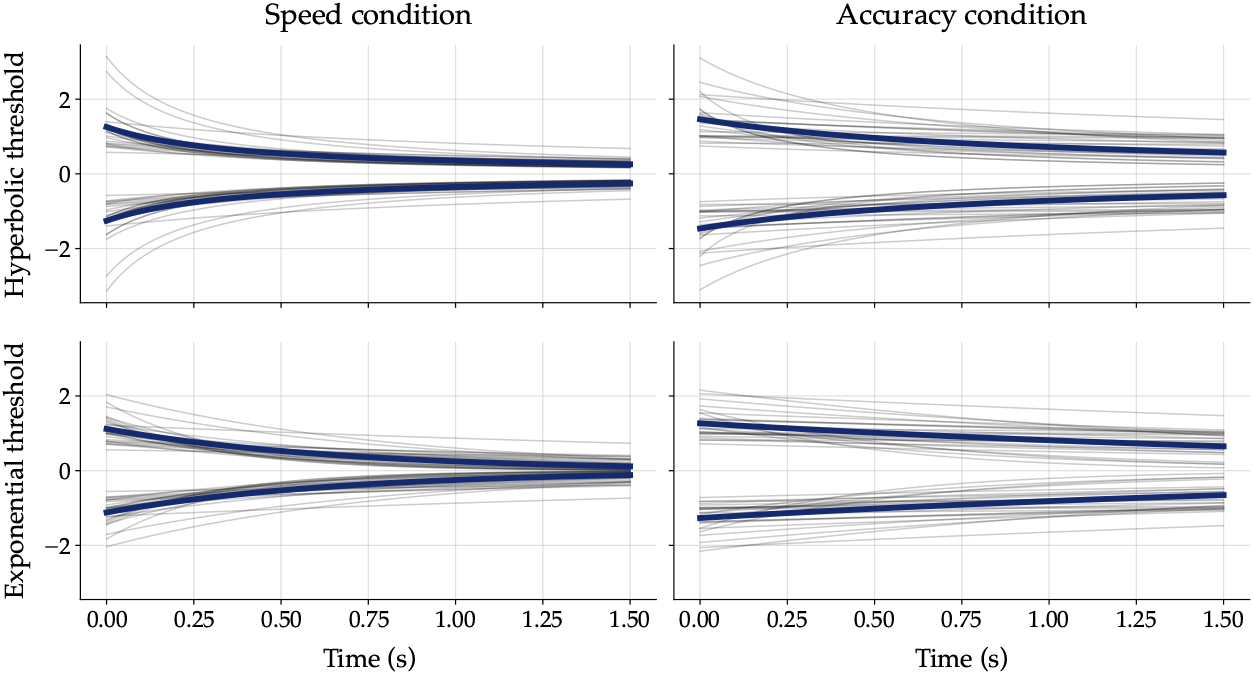
The estimated threshold dynamics from NDT-informed CT-DDMs for individuals in Study 1. The first row illustrates the estimated hyperbolic threshold dynamics, and the second row illustrates the estimated exponential threshold dynamics. The left column shows the threshold dynamics in the speed condition, and the right column shows the accuracy condition. Each gray line corresponds to a subject. The blue line corresponds to the average group level.

Figure 10 shows the predictions of the NDT-informed models for both the speed and accuracy conditions. The FT-DDM significantly overestimates the later RT quantile for correct and incorrect responses, in both speed and accuracy conditions. This overestimation is more pronounced in the speed condition. Moreover, FT-DDM underestimates the proportion of incorrect responses. In contrast to FT-DDM, which cannot accurately predict empirical data, particularly in the speed condition, the NDT-informed CT-DDMs exhibit high predictive accuracy.

**Figure 10.**
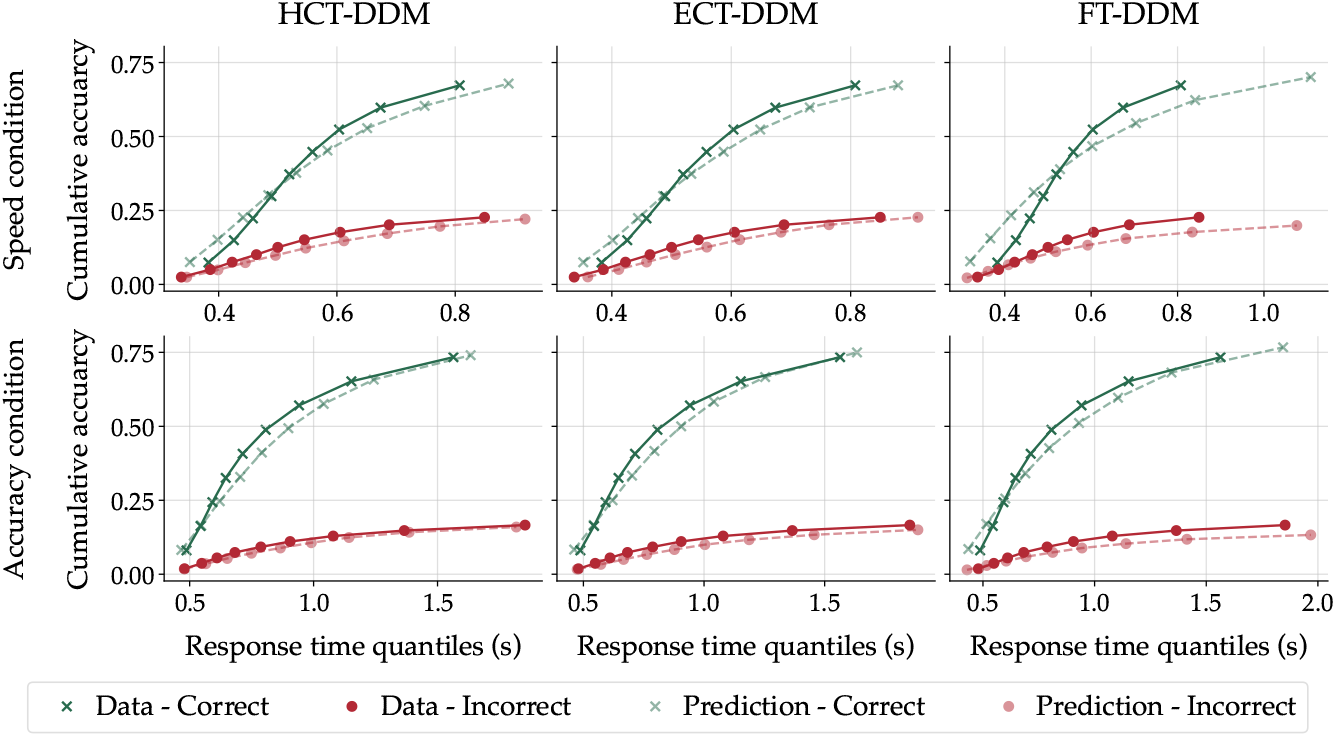
Prediction of NDT-informed models against empirical data for speed (top row) and accuracy (bottom row) conditions in Study 1. In each panel, the x-axis represents the response time quantiles in seconds, and the y-axis represents the cumulative choice proportion.

In sum, both quantitative and qualitative results revealed that the NDT-informed models perform better than the uninformed models in this study. In other words, constraining CT-DDM with non-decision time improved both the model fits and predictions. In addition to improving model fitting, the results obtained in this study also support the collapsing threshold against the fixed threshold. First, the mean estimated decay rates reported in Table 2 are significantly large. Second, all the considered CT-DDMs have a lower BIC and also better posterior prediction compared to the FT-DDM, which includes across-trial variability in drift rate (see the Appendix 7 for the results of FT-DDM). Third, the estimated threshold dynamics in Figure 9 indicate that most participants adopted a collapsing threshold. Therefore, considering all the results together, this study highlights the advantage of the NDT-informed modeling approach and supports CT-DDMs.

#### Study 2: Boehm et al. (2014) dataset

The dataset for the second case study is taken from ***Boehm et al. (2014***). In this experiment, twenty-five participants performed a random-dot motion task with a speed-accuracy trade-off manipulation. Before each trial, a cue (i.e., “AC” to emphasize accuracy in the subsequent trial or “SP” to emphasize speed in the following trial) was presented to inform participants how they should decide in the next trial. On each trial, participants viewed a cloud of moving dots for 1.5 s, of which a subset moved coherently in one direction while the rest moved randomly, and indicated the direction of coherent motion in a two-alternative forced-choice task. During the experiment, the coherency level (proportion of coherently moving dots) was fixed and determined in the practice block, ensuring that individuals achieved approximately 80% accuracy.

To model this dataset, we assumed that the drift rate remains fixed throughout the experiment. However, we allowed the threshold-setting and non-decision time to vary across speed and accuracy conditions.^12^ Similar to Study 1, we fitted HCT-DDM and ECT-DDM in two NDT-informed and uninformed versions, plus an FT-DDM, which includes across-trial variability in drift rate on empirical data. The non-decision time measurements were again obtained from a standard cumulative HMP fit (as reported in Section 4.4 of ***Weindel et al., 2024***). The HMP solution identified three events in the speed-focused condition and four events in the accuracy-focused condition. The sequence and timing of these events are presented in Figure 11. The non-decision time was taken as the difference between the RT and the duration of the second-to-last stage.

**Figure 11.**
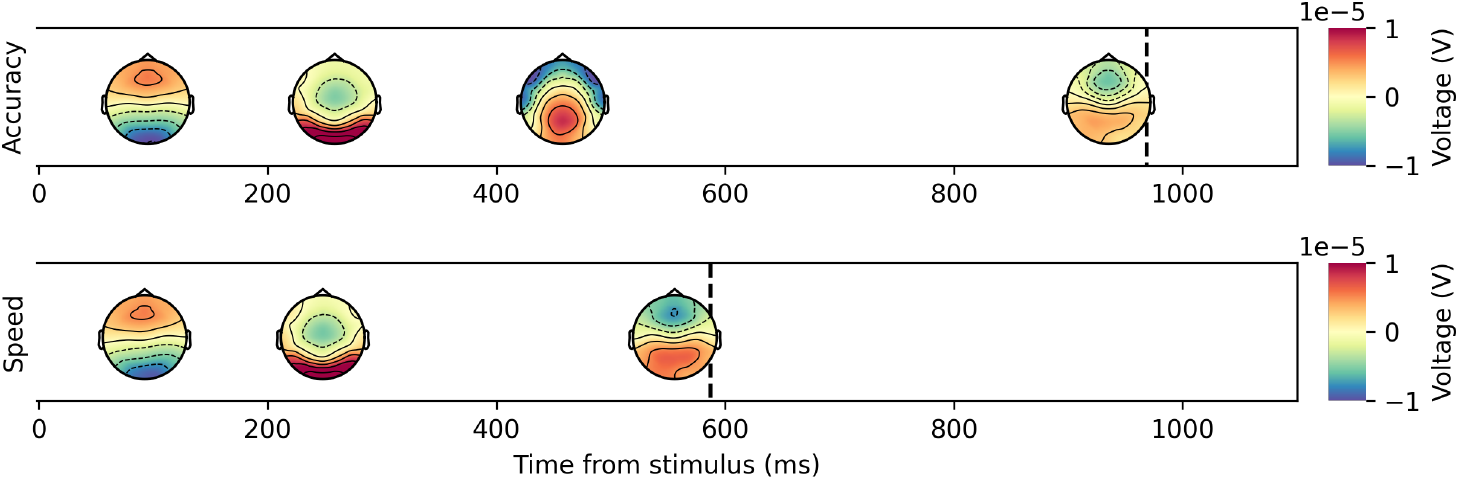
The event sequence and timing of each event estimated by the HMP method for Accuracy (top) and Speed (bottom) conditions averaged over all participants (***Weindel et al., 2024***).

As in Study 1, quantitative results showed that an NDT-informed model is the best-performing. The estimated mean parameter values and the model comparison results are presented in Table 3. The BIC-based model comparison suggests that the best-fitting model is the NDT-informed ECT-DDM. Therefore, the main result of the first study, which was an improvement in model fit when we constrain non-decision time, was replicated in this study.

**Table 3.** The mean estimated parameters and goodness of fit results for Study 2.

| Model | $\delta$ | $\tau_{SP}$ | $\tau_{AC}$ | $\theta_{SP}$ | $\theta_{AC}$ | $\lambda_{SP}$ | $\lambda_{AC}$ |
| --- | --- | --- | --- | --- | --- | --- | --- |
| NDT-informed HCT-DDM | 0.51 | 0.27 | 0.31 | 0.76 | 1.99 | 1.31 | 2.29 |
| NDT-informed ECT-DDM | 0.52 | 0.27 | 0.31 | 0.73 | 1.58 | 0.81 | 0.90 |
| NDT-informed FT-DDM | 0.57 | 0.29 | 0.43 | 0.56 | 0.72 | — | — |
| Uninformed HCT-DDM | 0.51 | 0.28 | 0.28 | 0.77 | 2.27 | 1.06 | 2.58 |
| Uninformed ECT-DDM | 0.53 | 0.27 | 0.31 | 0.74 | 1.57 | 0.72 | 0.87 |
| | $\sigma_{z,SP}$ | $\sigma_{z,AC}$ | BIC<sub>Joint</sub> | BIC<sub>Behavior</sub> | | | |
| NDT-informed HCT-DDM | 0.35 | 0.56 | -622.96 | 4164.78 |  |  |  |
| NDT-informed ECT-DDM | 0.35 | 0.58 | **-669.01** | **4116.75** |  |  |  |
| NDT-informed FT-DDM | 0.34 | 0.34 | 34.40 | 4466.14 |  |  |  |
| Uninformed HCT-DDM | — | — | — | 4268.05 |  |  |  |
| Uninformed ECT-DDM | — | — | — | 4216.85 |  |  |  |

The obtained thresholds differed notably between speed and accuracy conditions, with both the starting level and the decay rate being lower in the speed condition. The estimated threshold dynamics for each participant, along with the mean group estimated threshold, are presented in Figure 12 (see Appendix 6 for the estimated collapsing threshold by uninformed models). Similar to Study 1, the estimated thresholds for individuals suggest that most participants adopt a collapsing threshold stopping rule. Also, participants in this study exhibited a lower starting threshold in the speed condition compared to the accuracy condition. However, in contrast to Study 1, the decay rate in the speed condition was lower. This reduction in the decay rate under the speed emphasis condition is due to the huge effect of speed-accuracy threshold manipulation on the starting threshold.

**Figure 12.**
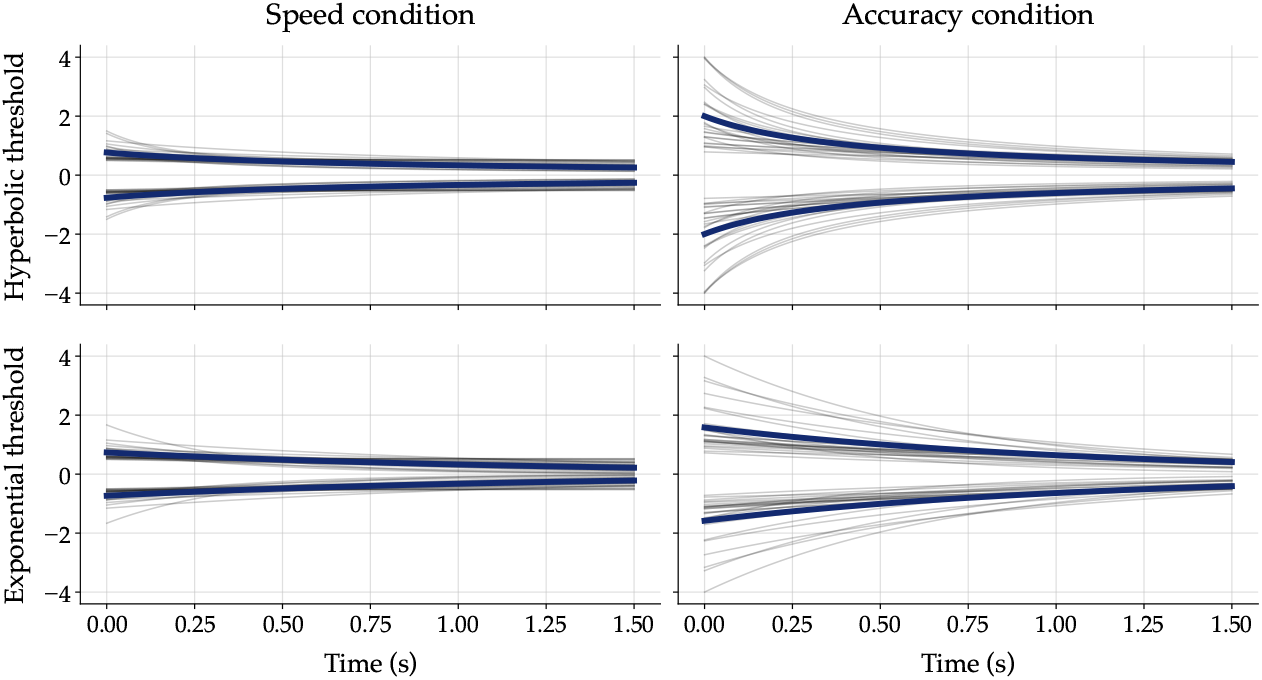
The estimated threshold dynamics from NDT-informed CT-DDMs for individuals in Study 2. The first row illustrates the estimated hyperbolic threshold dynamics, and the second row illustrates the estimated exponential threshold dynamics. The left column shows the threshold dynamics in the speed condition, and the right column shows the accuracy condition. Each gray line corresponds to a subject. The blue line corresponds to the average group level.

The NDT-informed models’ predictions are depicted in Figure 13. This figure shows that the FT-DDM overestimates the last RT quantiles for both correct and incorrect responses in both speed and accuracy conditions. However, the qualitative predictions of CT-DDMs align more closely with the empirical data. It is also worth noting that all the considered computational models misfit the incorrect responses in the speed condition. This misfit is to be linked to the presence of fast errors, specifically in the speed condition. Including the starting-point variability parameter in the model enables the model to predict fast errors. However, as the aim here was not merely to fit the data with the best possible model, but to test the NDT-informed modeling framework, we did not include starting-point variability in the model.

**Figure 13.**
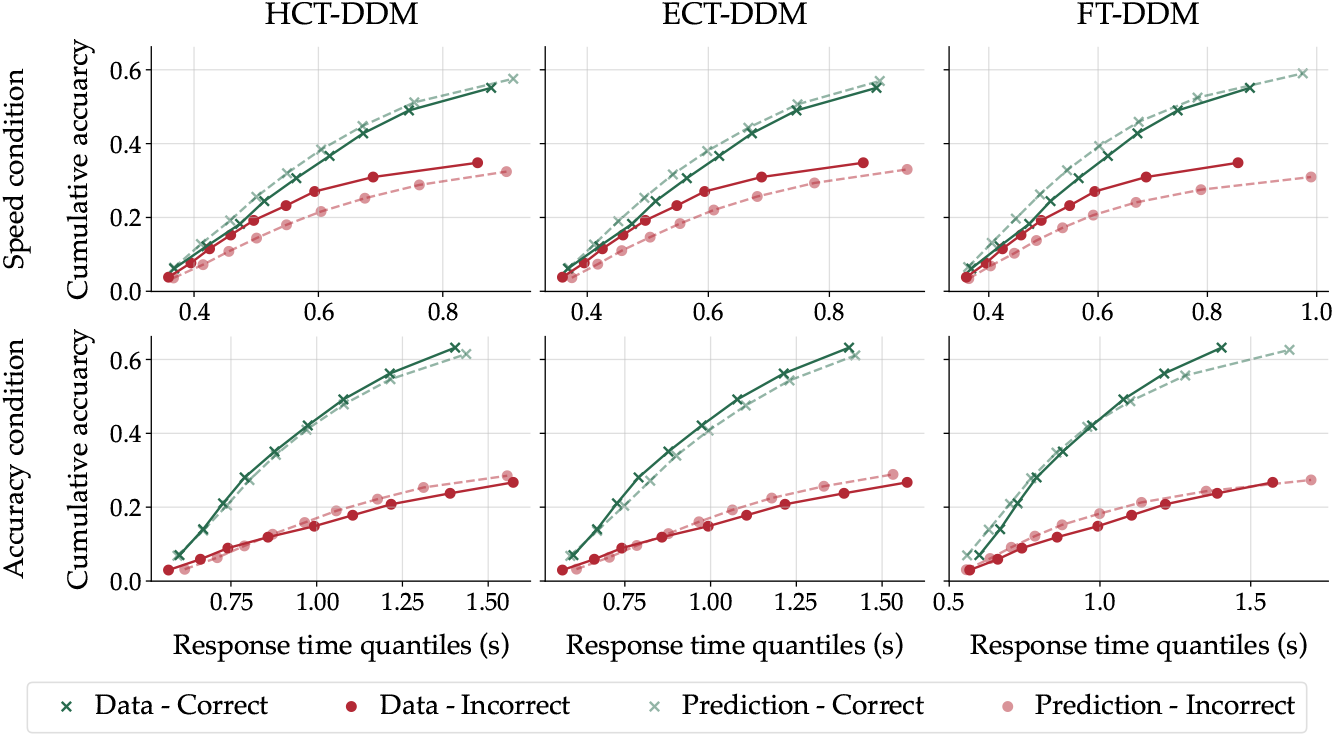
Prediction of NDT-informed models against empirical data for speed (top row) and accuracy (bottom row) conditions in Study 2. In each panel, the x-axis represents the response time quantiles in seconds, and the y-axis represents the cumulative choice proportion.

The results from this study replicated the main findings of the first study. The model comparison results showed that NDT-informed models fit the empirical data better than the uninformed models (in this study, NDT-informed ECT-DDM was the best-fitting model). Also, comparing these results with the results of FT-DDM reported in Appendix 7 revealed that the CT-DDMs better account for empirical data than the FT-DDM. This model comparison result, along with the high mean decay rate (reported in Table 3), supports the CT-DDMs against FT-DDMs.

### General discussion

Understanding whether people adjust their decision threshold over the time course of a single decision has been a central topic in cognitive science for decades. Despite the theoretical and empirical support for CT-DDMs, the main issue limiting their application in individual differences studies is the poor parameter recovery of these models. The present study addressed a long-standing question in computational cognitive neuroscience by proposing a diffusion modeling approach for the reliable estimation of time-dependent parameters. Specifically, we introduced a joint modeling framework that constrains non-decision time using additional observations extracted from neural signals. Simulation results demonstrated that incorporating non-decision time information significantly enhances the parameter recovery of collapsing threshold dynamics, making previously unrecoverable parameters recoverable. To test the robustness of this finding, the simulation results were replicated using two theoretically motivated forms of threshold dynamics: hyperbolic and exponential. While the precision of parameter estimation varied between these functional forms, the improvement in estimation was consistent across both models. That is, regardless of the threshold functional form, using noisy non-decision time measurements to inform the model consistently improved the reliability of parameter estimation. Importantly, the exponential collapsing threshold exhibited substantially better parameter recovery and superior model recovery performance, suggesting that this specification may be preferable in future cognitive modeling applications. Furthermore, the simulations showed that even under conditions of high noise in the non-decision time measurements, the parameter recovery of the NDT-informed model remained superior to that of the uninformed model. Additionally, the results of the model recovery simulation study showed that the NDT-informed modeling approach can reliably be used to infer the underlying evidence accumulation process. The model recovery study revealed that incorporating the estimated decay rate into model classification substantially increases model recovery precision.

In addition to the simulation studies, we reanalyzed two datasets on perceptual decision-making to demonstrate the feasibility and applicability of the proposed method in real empirical settings. In both studies, EEG recordings were available, and we used the HMP estimates from neural signals presented in the study by ***Weindel et al. (2024***) as measurements of non-decision time on a trial-by-trial basis. The cognitive modeling results revealed three main findings. First, informing CT-DDMs with non-decision time measurement improved model fit compared to their uninformed counterparts. This improvement likely reflects more accurate parameter estimation enabled by the additional information. In other words, constraining the non-decision time using neural measurements led the optimizer to estimate the CT-DDM parameter more accurately and, as a result, improve the fit to empirical data. Second, across both datasets, CT-DDMs outperformed FT-DDM in terms of model fit, regardless of whether the models were informed by non-decision time (see Appendix 7). This result, along with the better qualitative predictions of CT-DDMs, supports the notion that individuals adjust their decision thresholds dynamically within a single decision. Third, the estimated mean decay rate (*λ*) was notably high in both datasets, especially in the NDT-informed CT-DDMs — further supporting the presence of threshold collapse, at least in the analyzed datasets. Importantly, we found the evidence in two different types of experimental paradigms. In the second study, the information available to the participant for making their decision changed over the course of a single trial. However, in the first study, the stimulus remained static throughout the trial, and the amount of information remained constant. Most studies that have found evidence for urgency have employed information-changing paradigms (***Trueblood et al., 2021***; ***Evans et al., 2020a***; ***Evans and Hawkins, 2019***; ***Khodadadi et al., 2017***; ***Gluth et al., 2012, 2013***), and the support for collapsing the threshold in experiments with static stimuli is limited (see ***Olschewski et al., 2025***).

The computational modeling results contribute to the literature by providing empirical support for collapsing threshold models. The debate on whether people become more urgent as time passes is still ongoing. Although there are some empirical and theoretical supports for CT-DDMs, some studies based on quantitative model comparison across several experiments did not find strong support for collapsing threshold models (***Voskuilen et al., 2016***; ***Smith and Ratcliff, 2022***; ***Milosavljevic et al., 2010***; ***Karşılar et al., 2014***) or found that the collapsing threshold only happens in some specific circumstances (***Hawkins et al., 2015***). One reason for the mixed findings in the literature is the unreliability of parameter estimation in CT-DDMs (***Evans et al., 2020b***; ***Murrow and Holmes, 2024a***), along with high model mimicry in diffusion models (***Khodadadi and Townsend, 2015***), particularly when key parameters, such as the drift rate, vary across trials. However, the simulation results showed that based on NDT-informed diffusion modeling, we can reliably estimate the model’s parameters and infer the underlying evidence accumulation process. Thus, the proposed NDT-informed diffusion modeling approach can provide a stronger conclusion.

In addition to evidence supporting CT-DDMs, the computational modeling results also revealed that different types of speed-accuracy trade-off manipulations have distinct effects on the dynamics of the decision threshold. In Study 1, participants received verbal instructions emphasizing speed. This manipulation led to a faster decay rate and a decrease in the starting threshold. However, in Study 2, where participants faced a clear response deadline, the manipulation of speed caused a decrease in the decay rate. This decrease is mostly due to the large decrease in the starting threshold caused by speed emphasis in study 2. Task- and instruction-dependent effects of speed-accuracy trade-off manipulation have already been reported in several studies (e.g., ***Katsimpokis et al., 2020***; ***Evans et al., 2019***), and our results are consistent with this pattern.

One promising avenue for future research is the investigation of individual differences in the shape of threshold dynamics. In the present work, we examined two theoretically motivated forms — hyperbolic and exponential — and fitted them at the individual level. However, recent evidence suggests that threshold dynamics may vary substantially across individuals (e.g., ***Kira et al., 2025***). Non-parametric approaches have been proposed to recover the full threshold trajectory without assuming a specific functional form (e.g., ***Kira et al., 2025***; ***Fudenberg et al., 2020***). Findings from ***Kira et al. (2025***) indicate marked heterogeneity across participants. Understanding such variability could provide critical insights into why choice behavior sometimes deviates from optimality. A limitation of current non-parametric approaches, however, is that they require a vast number of trials per participant, making them impractical for studies with larger sample sizes. Combining NDT-informed diffusion modeling with non-parametric methods may help alleviate this limitation by reducing the trial requirements, thereby making individual differences studies on threshold dynamics more accessible to empirical investigation.

#### Limitation

The collapsing threshold dynamics considered in this work (e.g., exponential and hyperbolic) impose a monotonically decreasing threshold over time. Although these dynamics are theoretically well motivated (***Fudenberg et al., 2018***; ***Frazier and Yu, 2007***) and have been employed in several previous studies (e.g., ***Olschewski et al., 2025***; ***Milosavljevic et al., 2010***; ***Voskuilen et al., 2016***), some research has proposed delayed-collapsing threshold models (e.g., ***Diederich and Oswald, 2016***), in which the onset of threshold collapse does not coincide with the onset of evidence accumulation. Estimating such delayed-collapsing models may require more than simply constraining non-decision time, as the onset of threshold collapse must also be identified. A promising approach for addressing this challenge is the HMP method, which may provide additional temporal information about distinct cognitive processing stages. In particular, HMP may allow the onset of threshold collapse to be estimated as a separate cognitive stage. Future research should therefore investigate the estimation of multi-stage evidence accumulation models (e.g., ***Diederich and Oswald, 2016***; ***Diederich and Colonius, 2021***) within the HMP framework.

#### Distributional assumption on non-decision time measurement

Although we assumed that non-decision time measurements follow a log-normal distribution, alternative distributional assumptions are also plausible. In particular, some studies have modeled non-decision time using symmetric distributions such as the normal distribution (e.g., ***Ghaderi-Kangavari et al., 2023***; ***Kira et al., 2025***). In Appendix 8, we formalize the joint NDT-informed framework under the assumption that non-decision time measurements are normally distributed and report the corresponding parameter recovery results. The findings show that the improvement in recovering collapsing threshold parameters persists under this alternative distributional assumption. Although minor quantitative differences emerge across distributional specifications, the enhanced reliability of threshold parameter estimation is not confined to a specific assumption and generalizes across different distributional forms.

#### Would NDT-informed modeling lead to a different conclusion?

It is important to note that the NDT-informed diffusion modeling approach can sometimes yield conclusions that differ from those obtained using traditional modeling methods. For example, in Study 1, the results of the uninformed ECT-DDM suggest that the speed–accuracy manipulation does not affect the starting threshold. In contrast, both the NDT-informed ECT-DDM and NDT-informed HCT-DDM indicate that the speed–accuracy trade-off manipulation does influence the starting threshold (see Table 2). This discrepancy arises from unreliable parameter estimation in the uninformed CT-DDMs. However, the simulation results and sensitivity analyses reported in this paper demonstrate that the parameter estimates provided by the NDT-informed diffusion modeling framework are substantially more reliable. Therefore, any parameter-based inference concerning the underlying mechanisms of decision-making, the effects of experimental manipulations, or individual and group differences should be drawn using the NDT-informed diffusion modeling framework.

#### Generalization to other computational models

Although the primary focus of the present study was on two-alternative choice tasks, understanding urgency is also crucial in the context of multi-alternative and multi-attribute decisions (e.g., ***Tajima et al., 2019***; ***Gluth et al., 2026***; ***Busemeyer et al., 2019***), as well as in decisions involving a continuous range of options (e.g., circles, lines, or arcs; ***Hadian Rasanan et al., 2025, 2026, 2024a***). To examine the generalizability of the proposed method, we conducted a parallel parameter recovery study using the *n*-dimensional hyper-spherical diffusion model (***Smith and Corbett, 2019***; ***Smith, 2016***; ***Hadian Rasanan et al., 2026***). This model is suitable for modeling multi-alternative/multi-attribute choices (***Kvam, 2019a***; ***Smith and Corbett, 2019***; ***Smith, 2019***) as well as decision-making tasks with continuous responses (***Kvam, 2019b***; ***Smith et al., 2023, 2020***; ***Hadian Rasanan et al., 2026***). The results of this simulation study are presented in Appendix 9. Consistent with the findings from the one-dimensional DDM, the results show that informing the *n*-dimensional diffusion model with non-decision time measurements significantly improves the precision of threshold dynamics estimation (see Appendix 9). These findings suggest that the improvement in parameter recovery of the time-dependent threshold is not limited to binary choice tasks but also extends to the more general *n*-dimensional hyper-spherical diffusion model. This makes the proposed method applicable to a broader range of decision-making paradigms involving multi-alternative, multi-attribute, or continuous response decision tasks.

The proposed NDT-informed diffusion modeling approach can also be applied to parameter estimation in urgency gating models. Unlike CT-DDMs, urgency-gating models assume a fixed decision threshold, but allow the incoming evidence to scale as time progresses (***Trueblood et al., 2021***; ***Cisek et al., 2009***). However, similar to CT-DDMs, urgency-gating models also suffer from poor parameter recovery, as reported by ***Evans et al. (2020b***). Thus, the issue of parameter unreliability extends to these models. Recently, ***Smith and Ratcliff*** (***2022***) formalized the connection between CT-DDMs and urgency gating models, demonstrating that the multiplicative urgency gating model is mathematically equivalent to a hyperbolic collapsing threshold diffusion model. In other words, urgency gating models and hyperbolic CT-DDMs yield identical behavioral predictions, and their parameters can be transformed into one another. Consequently, the findings from our simulation studies on hyperbolic collapsing threshold dynamics also apply to urgency gating models (***Trueblood et al., 2021***; ***Cisek et al., 2009***).

Another way to implement urgency in the decision process is to consider an independent accumulator for timing. The time-based racing diffusion model (***Hawkins and Heathcote, 2021***) considers independent accumulators corresponding to evidence for available options, as well as an additional time-based accumulator for timing. This time-based accumulator behaves like an inner clock, and when it exceeds the threshold before the other evidence accumulators, an option will be randomly selected. Similar to the CT-DDM, this model can also generate Gaussian-like distributions of RTs and shows a good fit on empirical data. This model can also predict faster error responses than the correct response, the pattern that is observed in the speed condition of Study 2. However, the authors reported poor parameter recovery for the onset of the timing process (***Hawkins and Heathcote, 2021***). Thus, the reliability issue here specifically concerns the estimation of the shift parameter of the timing accumulator. Informing the model with external estimates of non-decision time might, therefore, improve parameter recovery in this model as well.

More generally, collapsing thresholds are not the only evidence accumulation mechanisms that suffer from reliability issues. Poor parameter recovery has also been reported for time-dependent drift rate models, such as diffusion models of conflict or diffusion models with leakage (***White et al., 2018***; ***Evans et al., 2020b***). Although several studies have attempted to improve parameter recovery in these models (e.g., ***Hübner and Pelzer, 2020***; ***Miletić et al., 2017***), the problem remains unresolved. A promising direction for enhancing parameter estimation in such models is to constrain the drift rate dynamics using neural data. In particular, several studies have shown that Centro-parietal positivity (CPP) in EEG signals is closely linked to the process of evidence accumulation (e.g., ***O’Connell et al., 2012***; ***Steinemann et al., 2018***; ***Kohl et al., 2020***; ***Kelly et al., 2021***; ***Grogan et al., 2025***). Furthermore, a recent application of HMP has shown that single-trial estimates of this potential can be extracted with reasonable accuracy (***Weindel et al., 2025***). Thus, informing time-dependent drift rate models with CPP dynamics may improve the identifiability and reliability of their parameter estimates.

#### Does only non-decision time improve parameter estimation?

In this work, we focused on constraining the non-decision time and provided both theoretical justification and simulation evidence demonstrating that constraining non-decision time in the diffusion model can enhance the reliability of parameter estimation in CT-DDMs. However, it remains an open question whether constraining other parameters — such as the drift rate — can similarly improve parameter estimation in CT-DDMs. To address this, it is essential to note that, from a mathematical standpoint, assuming a fixed diffusion coefficient, there is no trade-off between drift rate and threshold parameters (***Nunez et al., 2025***; ***Ratcliff, 1978***). Therefore, constraining the drift rate using neural signals would not improve the estimation of collapsing threshold parameters.

Another neurophysiological signal that may aid in improving parameter estimation of CT-DDMs is pupil dilation. Several studies have demonstrated a correlation between pupil dilation and decision threshold (e.g., ***Murphy et al., 2016***; ***Cavanagh et al., 2014***), suggesting that the pupil dilation time series may contain valuable information about threshold dynamics. However, this approach presents certain challenges. Notably, accurately identifying the time window corresponding to the decision process using only eye-tracking data is difficult. To overcome this limitation, additional sources of information — such as EEG signals — may be necessary. As a result, directly constraining threshold dynamics is more complex than constraining non-decision time. Nevertheless, the potential advantages of incorporating pupil dilation into joint modeling approaches warrant further investigation in future research.

#### Bias in non-decision time measurement

One of the main assumptions of NDT-informed diffusion modeling is that the true non-decision time equals the mean of the non-decision time measurement distribution. However, this assumption may not always hold in certain situations. In other words, there might be a systematic bias (i.e., overestimation or underestimation) in the non-decision time measurements. To investigate the effect of such biases in non-decision time measurements on parameter estimation, we conducted a model misspecification study in which the true non-decision time parameter deviates from the mean of non-decision time measurements (see Appendix 5). The results of the model misspecification study showed that when we have an overestimation bias in non-decision time measurements, we observe an underestimation in the starting threshold and decay rate parameters (with high correlation with the true parameters). In contrast, when there is an underestimation bias in non-decision time measurements, we observe an overestimation in starting threshold and decay rate parameters.

#### Behavioral methods for estimating non-decision time

Although the focus of the current study was on extracting non-decision time from neural signal data using the HMP method, it is also worth noting that some alternative approaches do not require neural data. Historically, disentangling decision time and non-decision time has been a long-standing question in mathematical psychology and psychophysics (e.g., ***Green and Luce, 1971***; ***Kohfeld et al., 1981***; ***Burbeck and Luce, 1982***; ***Smith, 1990***; ***Sheu and Ratcliff, 1995***; ***Verdonck and Tuerlinckx, 2016***; ***Bompas et al., 2025***). These attempts resulted in the development of convolutional-based methods. The core idea behind these methods is that the response time distribution can be represented by the convolution of the decision time distribution predicted by the diffusion model and the non-decision time distribution (***Smith, 1990***; ***Verdonck and Tuerlinckx, 2016***). Therefore, one way of using this relation is to deconvolve the non-decision time distribution from the observed RT distribution, resulting in the decision time distribution (***Smith, 1990***). Thus, running a simple one-choice reaction task (i.e., responding as soon as the stimulus is presented on the screen) with identical stimuli, which are also presented in the main task, provides an estimation of the non-decision distribution. In other words, response time in such one-choice reaction tasks includes only the perceptual encoding and motor execution, excluding the evidence accumulation process, and approximates the non-decision time in the main task. Deconvolution of the non-decision time distribution obtained from the one-choice reaction task to the response time distribution of the main task provides an approximation of decision time in the main task. This approach requires running an additional task and also has some practical issues (***Sheu and Ratcliff, 1995***).

To address these issues, ***Verdonck and Tuerlinckx*** (***2016***) proposed an alternative convolution-based approach that can estimate the distribution of non-decision time entirely based on observed data in a single experiment. They estimate the non-decision time distribution by convolving the observed RT distribution with the predicted distribution from the diffusion model in different conditions and minimizing the distance between the resulting distributions. Their results showed that their proposed method can accurately estimate the distribution of non-decision times and cancel out the effect of non-decision time on RT. Recently, ***Kira et al. (2025***) have employed a similar approach to cancel out the impact of non-decision time on RT and then estimate the entire threshold dynamics using a non-parametric approach.

## Conclusion

The question of whether people become more urgent as time passes is one of the long-standing questions in cognitive psychology. In this work, we introduced an NDT-informed diffusion modeling framework that significantly improves the reliability of parameter estimation in CT-DDM. Therefore, this approach enables researchers to use parameter values in their inference about the underlying cognitive mechanism of the decision process and to use CT-DDM as a measurement tool to investigate individual differences. Based on the proposed NDT-informed diffusion modeling framework, we leveraged EEG estimates of non-decision time and reanalyzed two empirical datasets. We found evidence for CT-DDMs across two studies, which implies that people become more urgent as time passes.

## Acknowledgments

The authors would like to thank Nathan J. Evans, Sebastian Gluth, and Amin Ghaderi-Kangavari for their insightful comments on the early version of this work.

## Declarations

### Code and data availability

All codes and data supporting this paper are publicly available on: https://github.com/AmirHoseinHadian/CTDM_recovery

## Appendix 1

### Trade-off between non-decision time and collapsing threshold recovery

To illustrate how imprecise estimation of non-decision time affects the recovery of collapsing threshold parameters (i.e., starting threshold and decay rate), we conducted an additional simulation study. In this study, data was generated from an uninformed CT-DDM, after which model parameters were estimated under the assumption that non-decision time was fixed rather than estimated freely. To show how imprecise estimation of non-decision time can distort threshold estimation, the non-decision time was either held fixed or contaminated with noise. Using this scenario and modulating the noise level, we can understand how and on what scale noisy estimation of non-decision time affects threshold estimation.

To simulate data, first, we generated sample parameters from the following uniform distributions:

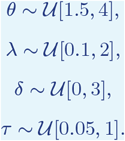

After sampling the parameters, we generated response time and choice for 500 trials using the drawn parameters. Then, we estimated parameters using the simulated data. But during the estimation procedure, the non-decision time parameter was treated as known and held constant. In other words, we only estimated *θ, λ*, and *δ*. The fixed non-decision time value was either exact or contaminated with uniform noise at five levels (2%, 4%, 6%, 8%, or 10%). The contaminated non-decision time is obtained by *τ*_contaminated_ = (1 + *ε*) *× τ*, where *ε* ∼ 𝒰 [−*n*, +*n*], and *n* is either 0.02, 0.04, 0.06, or 0.1.

Figure 1 illustrates the effect of noisy non-decision time estimation on the recovery of other model parameters in both exponential and hyperbolic CT-DDMs. As shown in this figure, increasing noise in the non-decision time estimates substantially degrades the estimation of the starting threshold (*θ*) and decay rate (*λ*) parameters. Specifically, the root mean square error (RMSE) for these parameters increases as the noise level in non-decision time increases. In addition, both the correlation coefficient (*ρ*) and the *R*^2^ value for these parameters decrease with increasing non-decision time noise. In contrast, these metrics for the drift rate remain largely unchanged, indicating that drift rate estimation is relatively unaffected by imprecise non-decision time estimates. These simulation results are consistent with the theoretical arguments presented in the main text and confirm the existence of a trade-off between non-decision time and collapsing threshold parameters.

**Appendix 1—figure 1.**
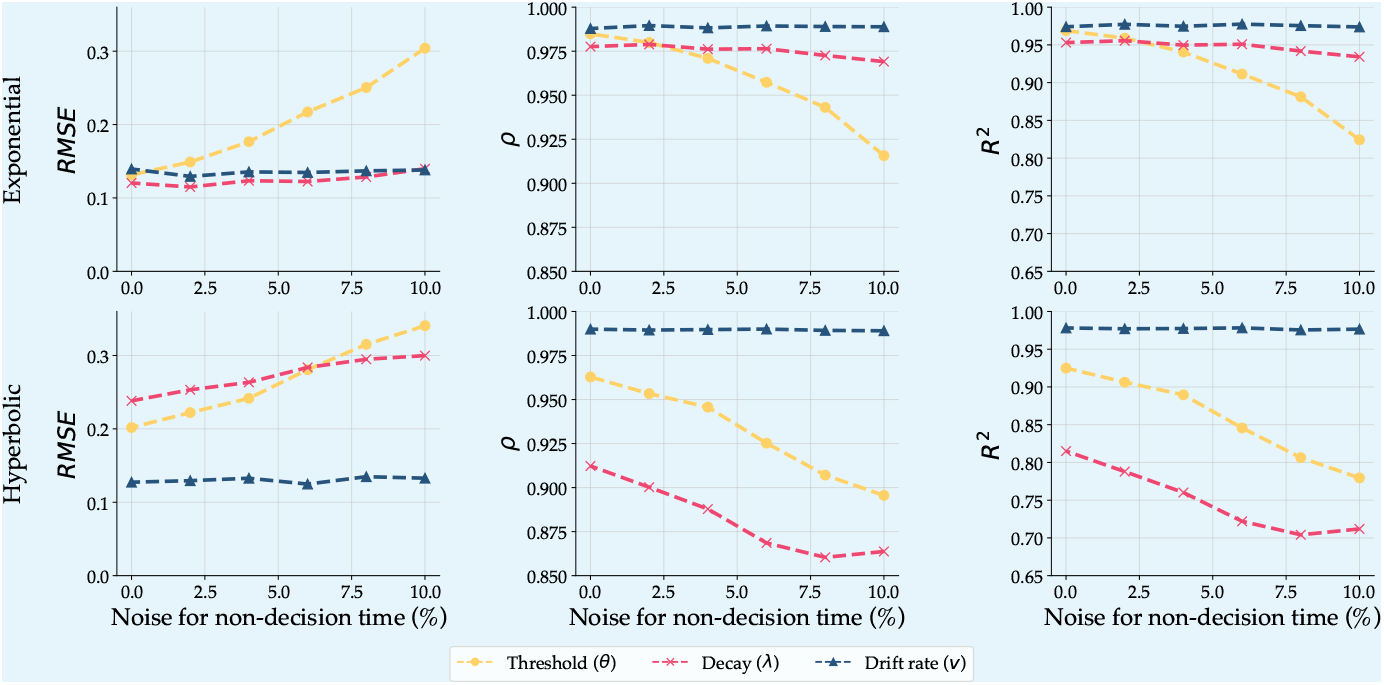
Illustration of the imprecise estimation of non-decision time on other parameters of the CT-DDM.

## Appendix 2

### Model fitting using the integral equation method

Since the diffusion models with time-dependent thresholds do not have an exact likelihood function, we need to approximate it using a numerical method. The integral equation method provides a reliable numerical procedure for approximating the first-passage time distribution of diffusion models with time-dependent parameters (***Smith, 2000***; ***Smith and Ratcliff, 2022***). This method was first introduced by ***Buonocore et al. (1987***, 1990) and then employed in psychological research frequently (e.g., ***Smith, 2000***; ***Voskuilen et al., 2016***; ***Smith and Ratcliff, 2022***; ***Evans et al., 2020a***; ***Hadian Rasanan et al., 2025***; ***Smith, 2023***; ***Zhang et al., 2014***). In this method, the first-passage time distributions of upper and lower boundaries are obtained by solving the following system of linear integral equations (in mathematics, this equation is known as the Volterra integral equation of the second kind, ***Wazwaz, 2011***):

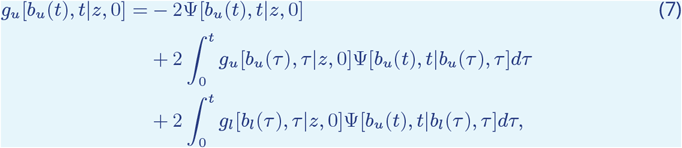

and

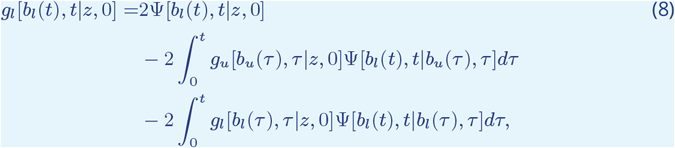

where *g*_*u*_[*b*_*u*_(*t*), *t*|*z*, 0] and *g*_*l*_[*b*_*l*_(*t*), *t*|*z*, 0] represent the first-passage time distributions of crossing the upper and lower time-dependent thresholds, respectively. Also, Ψ is the kernel function and is defined as follows (***Giorno et al., 1989***; ***Richter et al., 2023***):

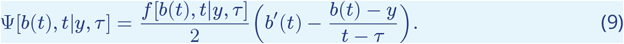

In this equation, *f* [*x, t*|*y, τ* ] represents the free transition density function. For the one-dimensional diffusion process with drift rate *δ*, the free transition density function is defined as follows (***Giorno et al., 1989***; ***Richter et al., 2023***):

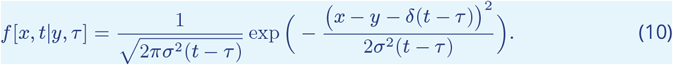

To approximate the solution of this integral equation, various numerical methods are available in the literature (see Chapter 3 in ***Wazwaz, 2011***). However, the left-point rectangular scheme is the most straightforward and efficient numerical approximation scheme, offering relatively high precision in likelihood approximation (***Richter et al., 2023***). So, by discretizing the temporal space using *N* equidistant points (i.e., 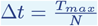, we considered *T*_*max*_ equal to the longest RT) the approximation of the first-passage time at the first time step is obtained as follows (***Buonocore et al., 1990***; ***Smith, 2000***):

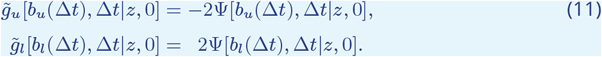

Also, for the next time steps (i.e., *t*_*i*_ = *i*Δ*t* for *i* = 2, 3, … ) we can use the following approximation scheme (***Buonocore et al., 1990***; ***Smith, 2000***):

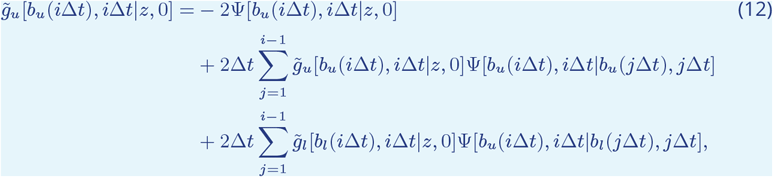

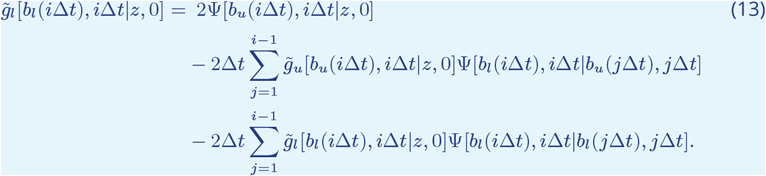

After approximating the first-passage time distributions corresponding to the upper and lower thresholds, the negative log-likelihood can be estimated using:

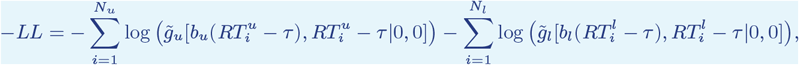

where 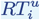 and 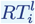 represent the response times ended at the upper threshold and lower threshold, respectively. Moreover, *N*_*u*_ is the number of trials in which the process stopped at the upper threshold, and similarly, *N*_*l*_ is the number of trials in which the process terminated at the lower threshold.

## Appendix 3

### Simulation results for linear collapsing threshold diffusion model

Although there are strong theoretical motivations for considering nonlinear collapsing threshold dynamics in diffusion models, linear CT-DDMs are frequently used in the literature. This widespread use is largely due to the relatively better parameter recovery of linear CT-DDMs compared to their nonlinear counterparts. As discussed earlier, ***Evans et al. (2020b***) conducted a systematic assessment of parameter recovery in CT-DDMs and found that the parameters of linear CT-DDMs can be reliably identified given a sufficiently large number of trials. To examine whether constraining non-decision time can further improve parameter recovery and reduce the number of trials required for reliable estimation, we conducted a simulation study comparing NDT-informed and uninformed linear CT-DDMs. In this study, we assumed a linearly collapsing decision threshold (i.e., *b*_*u*_(*t*) = −*b*_*l*_(*t*) = *θ* − *λ × t*). All other aspects of the simulation design were identical to those used in the simulation study reported in the main text.

Figure 1 shows the *R*^2^ values for the parameters of both uninformed and NDT-informed linear CT-DDMs across different numbers of trials. In addition, Figure 2 displays the relationship between the estimated parameters and the true generating parameters for the simulation with 500 trials. Consistent with the simulation results reported by ***Evans et al. (2020b***), the results for the uninformed linear CT-DDM indicate that the parameters become identifiable when a sufficiently large number of trials is available (i.e., more than 500 trials). However, as with nonlinear CT-DDMs, constraining the non-decision time substantially improves the estimation of collapsing-threshold parameters. Notably, the starting threshold parameter can be reliably identified with as few as 100 trials. Thus, although the parameters of the linear CT-DDM can be accurately estimated given enough data, constraining non-decision time markedly reduces the number of trials required for precise parameter estimation.

**Appendix 3—figure 1.**
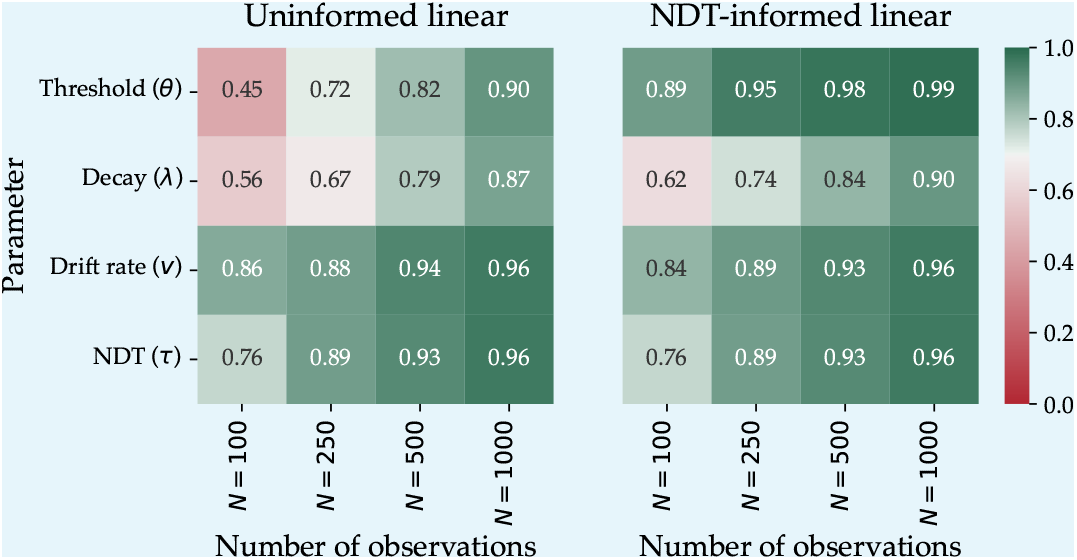
*R*^2^ values measuring the agreement between estimated and ground truth parameters in uninformed (left) and NDT-informed (right) linear collapsing threshold diffusion models as a function of the number of observations.

**Appendix 3—figure 2.**
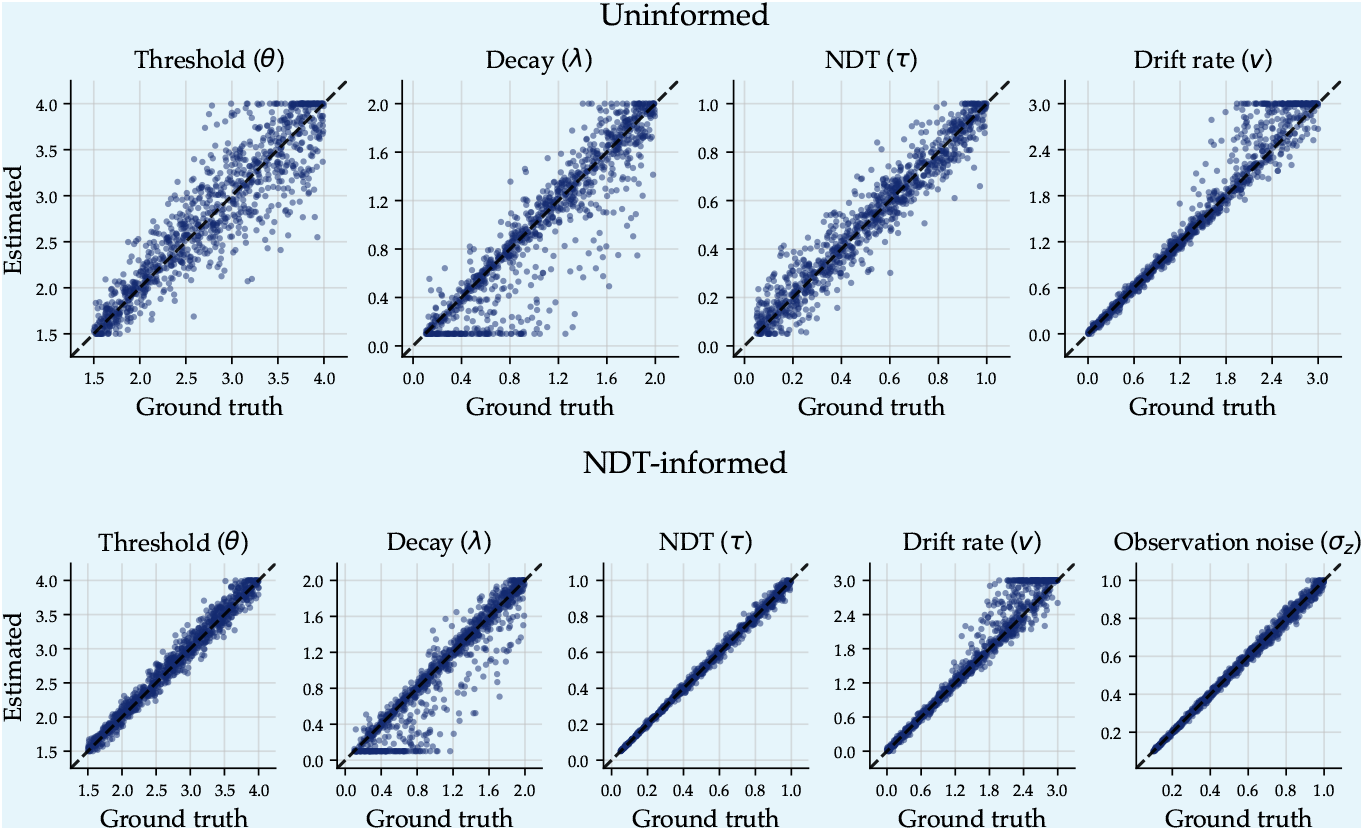
Estimated vs. true parameter values for two modeling approaches. The top row shows parameter recovery for the linear collapsing threshold diffusion model estimated without additional constraints (“Uninformed”), while the bottom row shows recovery for the joint model that incorporates additional non-decision time observations (“NDT informed”). Each point represents a recovered parameter estimate plotted against its corresponding true data-generating value. Dashed diagonal lines indicate perfect recovery.

## Appendix 4

### Sensitivity to estimation method

Sometimes, problems in parameter recovery are specific to a particular estimation method and may not occur with others. To ensure the robustness of our results, we conducted the same simulation study using an alternative estimation approach – Amortized Bayesian Inference (ABI; ***Radev et al., 2020***; ***Bürkner et al., 2023***). ABI is a modern inference technique that leverages neural networks to approximate posterior distributions efficiently. Instead of performing Bayesian inference from scratch for each new dataset, ABI “amortizes” the computational cost by learning a mapping from observed data to posterior distributions across many simulated datasets.

For this purpose, we utilized the BayesFlow Python library (***Radev et al., 2023b***), which provides neural network architectures tailored for ABI. Our inference pipeline consists of two main components: a set transformer that encodes the trial-wise data into maximally informative summary statistics, and an invertible neural network (the inference network) that takes these summaries — along with the generative parameters — and learns to approximate the posterior distribution. We trained four separate neural approximators, corresponding to each of the models examined: NDT-informed ECT-DDM, NDT-informed HCTDDM, Uninformed ECT-DDM, and Uninformed HCT-DDM. Each network was trained on 2 400 000 simulated datasets, with each dataset containing 500 trials and parameters sampled from prior distributions that matched those used in the main simulation study. Further implementation details, including network architectures and hyperparameters, are available in our publicly accessible GitHub repository: (https://github.com/AmirHoseinHadian/CTDM_recovery)

Figure 1 displays the *R*^2^ values for the threshold parameters (i.e., *θ* and *λ*) under both the no-constraint and joint models. Applying constraints on non-decision time significantly enhances parameter recovery accuracy, with the greatest improvement observed in the Hyperbolic collapsing threshold model. Compared to the integral equation estimation method, parameter recovery for the non-informed models was significantly better – again, especially for the Hyperbolic collapsing boundary. In contrast, *R*^2^ values for the NDT-informed parameters were comparable across both methods. For completeness, Figure 2 presents the full parameter recovery results for all four models. This figure illustrates the estimated parameters against the true generating parameters for each model.

**Appendix 4—figure 1.**
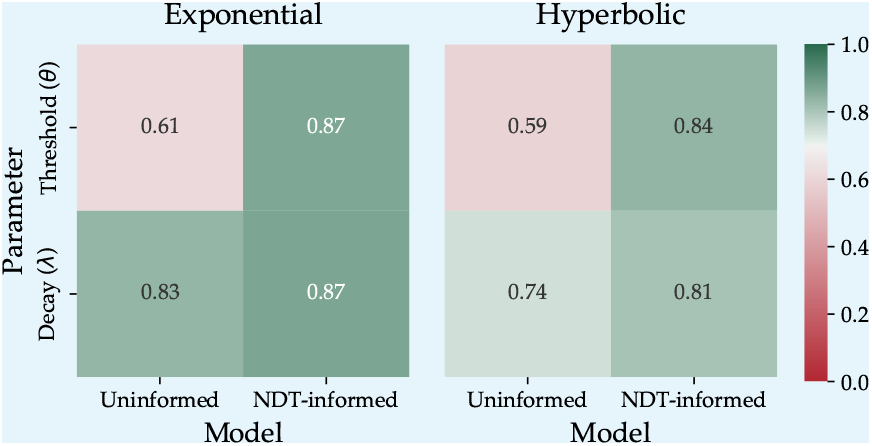
*R*^2^ values measuring the agreement between estimated and ground truth threshold parameters in exponential (left) and hyperbolic (right) collapsing boundary models.

**Appendix 4—figure 2.**
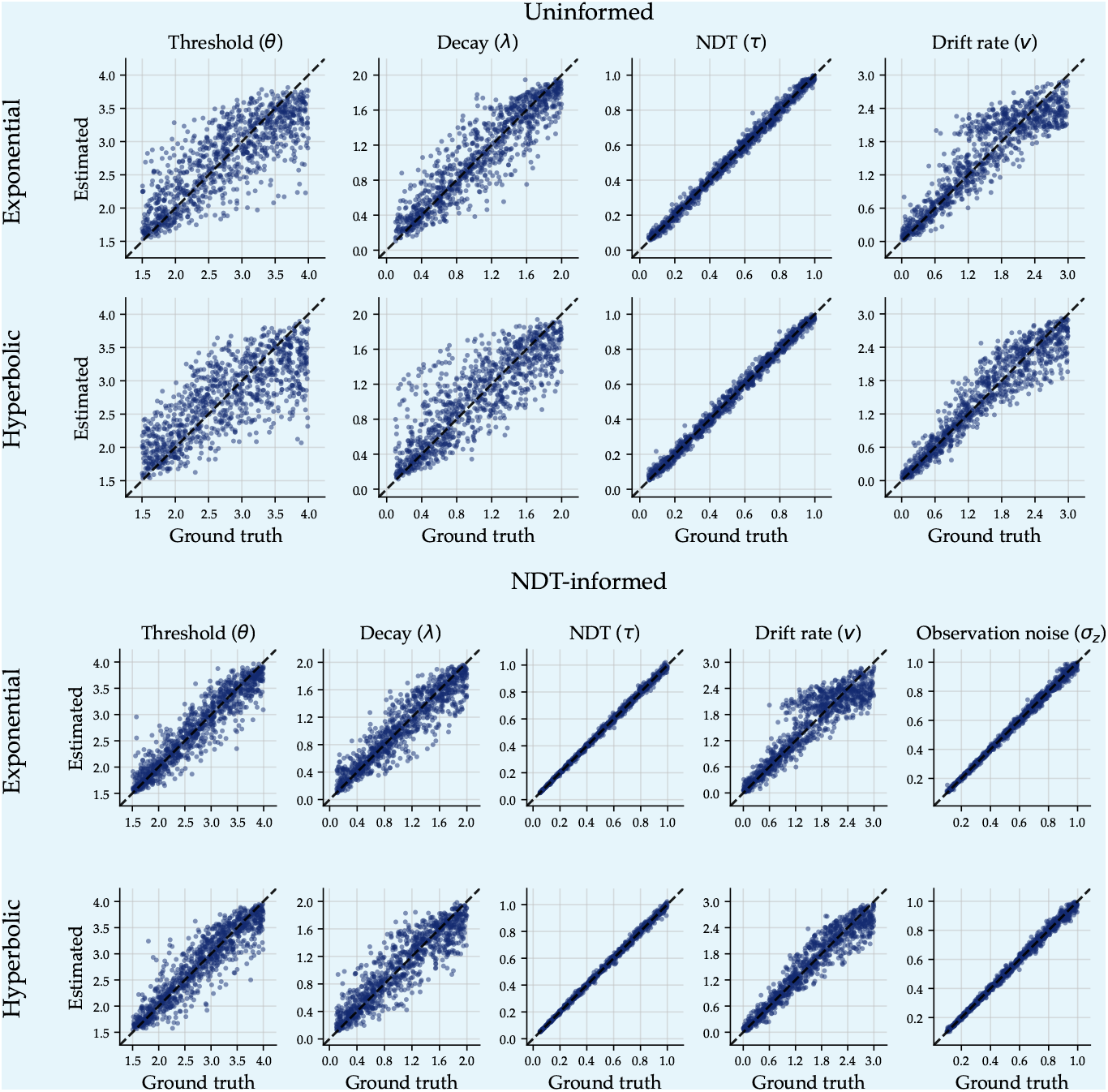
Estimated vs. true parameter values for two modeling approaches. The top two rows show parameter recovery for models estimated without additional constraints (“Uninformed”), while the bottom two rows show recovery for joint models that incorporate additional non-decision time observations (“NDT informed”). Each point represents a recovered parameter estimate plotted against its corresponding true data-generating value. Rows correspond to different functional forms (Exponential and Hyperbolic). Dashed diagonal lines indicate perfect recovery.

## Appendix 5

### Bias in non-decision time measurements

To investigate the impact of bias in non-decision time measurement on parameter estimation of NDT-informed CT-DDMs, we conducted a simulation study. The simulation setup for data generation was identical to the one considered in the main text, except that the mean of non-decision time measurements (*Z*_*n*_) distribution deviated from the true non-decision time parameter (*τ* ). To generate non-decision time measurements, we considered the following biased distribution:

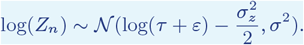

where *ε* represents the bias in the scale of milliseconds. We considered six levels of bias in measurements (i.e., −60, −40, −20, +20, +40, and +60 milliseconds). These values cover systematic bias in the range of 60 milliseconds underestimation to 60 milliseconds overestimation. This deviation is not the maximum deviation; rather, it represents a systematic bias in addition to the variability in the non-decision time measurements.

Figure 1 illustrates the effect of measurement bias on parameter estimation based on the *R*^2^ value. The results suggest that the threshold parameters are sensitive to measurement bias. This sensitivity is smaller for the exponential model. At first glance, this figure may suggest that the recovery of threshold parameters becomes increasingly unreliable as the bias increases. However, it is worth noting that these low *R*^2^ values result from bias in parameter estimation when there is a bias in non-decision time measurements.

**Appendix 5—figure 1.**
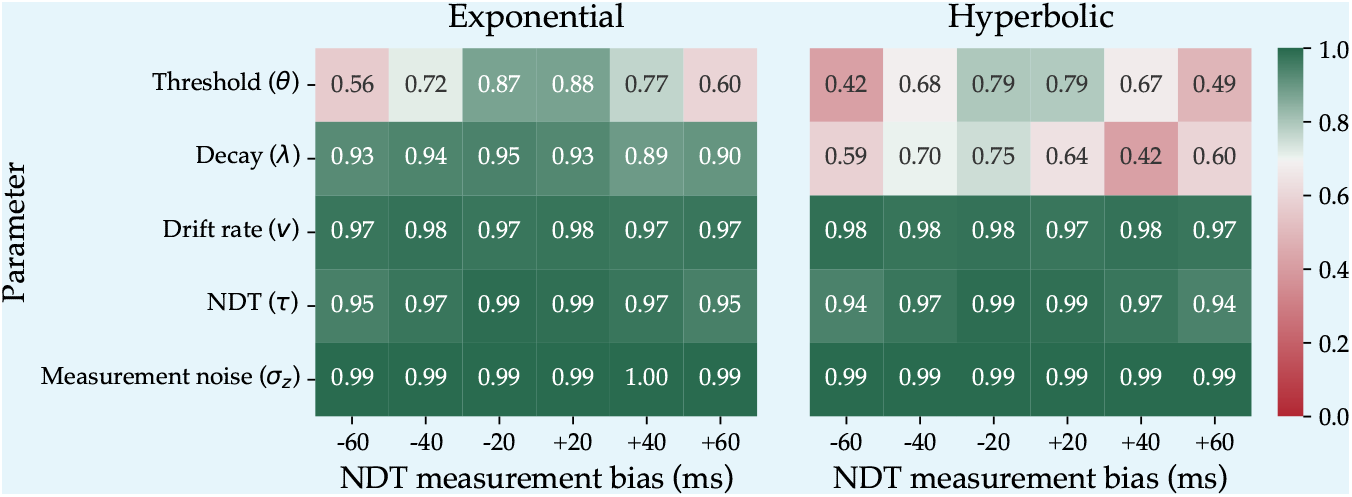
*R*^2^ values measuring the agreement between estimated and ground truth parameters in exponential (left) and hyperbolic (right) collapsing threshold diffusion models as a function of bias in non-decision time measurements.

To illustrate this, Figure 2 shows the correlation between the true parameters and the estimated parameters in the presence of bias in non-decision time measurement. The obtained correlation values are very high, suggesting that the poor *R*^2^ values reported in Figure 1 are mostly due to a bias in parameter estimation rather than variability or unreliability in the parameter estimation.

**Appendix 5—figure 2.**
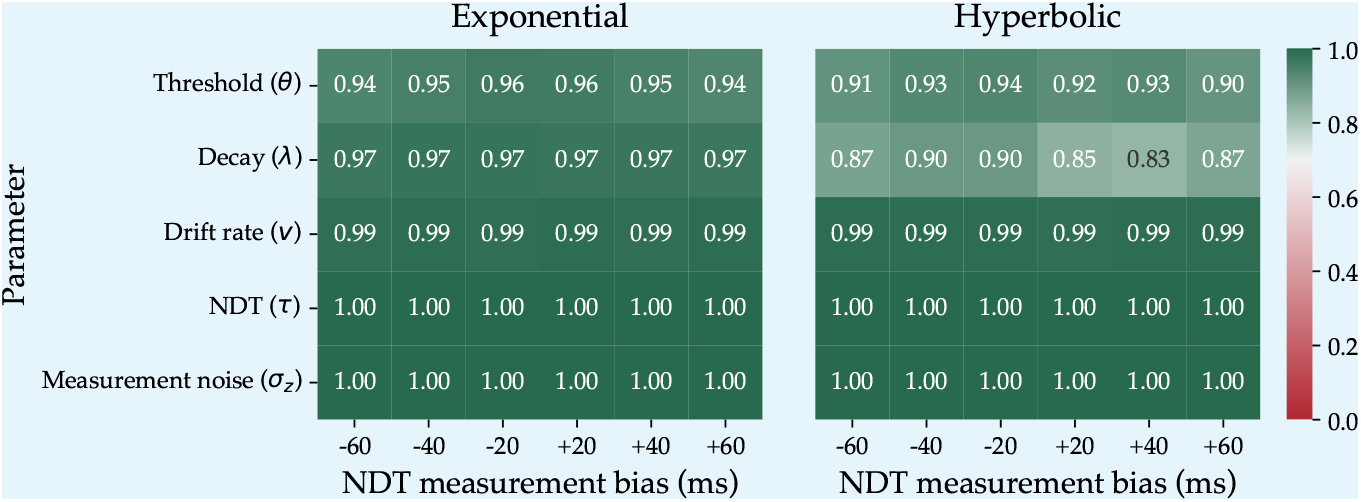
The correlation values measuring the agreement between estimated and ground truth parameters in exponential (left) and hyperbolic (right) collapsing threshold diffusion models as a function of bias in non-decision time measurements.

Figure 3 presents the effect of bias in non-decision time measurements on threshold parameters. Underestimation in non-decision time leads to overestimation in the starting threshold and decay rate. Conversely, overestimation in non-decision time leads to underestimation in both the starting threshold and non-decision time. The effect of bias is larger on the starting threshold than on the decay rate. Moreover, the exponential model was less sensitive to the bias in non-decision time measurements than the hyperbolic model.

**Appendix 5—figure 3.**
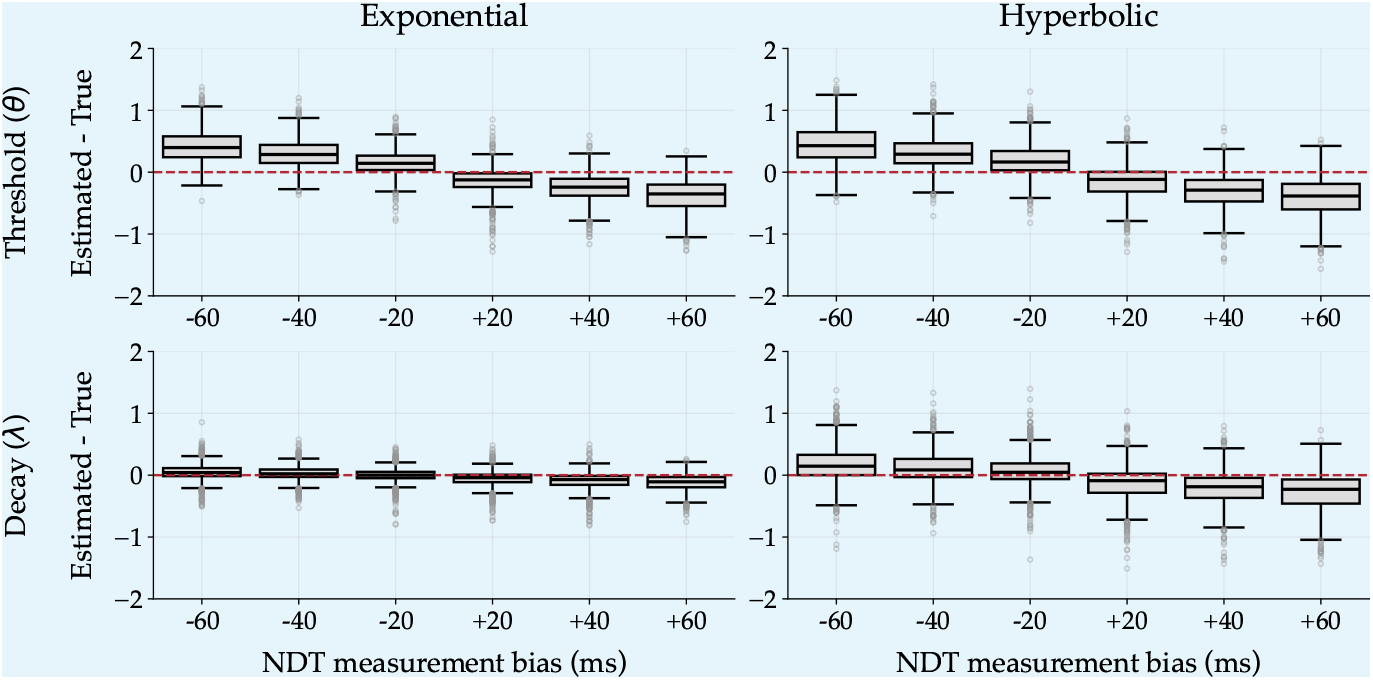
The effect of bias in non-decision time measurements on starting threshold (upper panels) and decay rate (lower panels) parameters for exponential (left panels) and hyperbolic (right panels) collapsing threshold diffusion models.

## Appendix 6

### Threshold estimation for uninformed CT-DDMs

This appendix presents the estimated threshold dynamics for uninformed CT-DDMs. Figure 1 and Figure 2 illustrate the estimated thresholds for each individual in study 1 and study 2, respectively. Similar to the obtained results for NDT-informed CT-DDMs, here we also observed that the decision threshold for most participants collapses largely over time.

**Appendix 6—figure 1.**
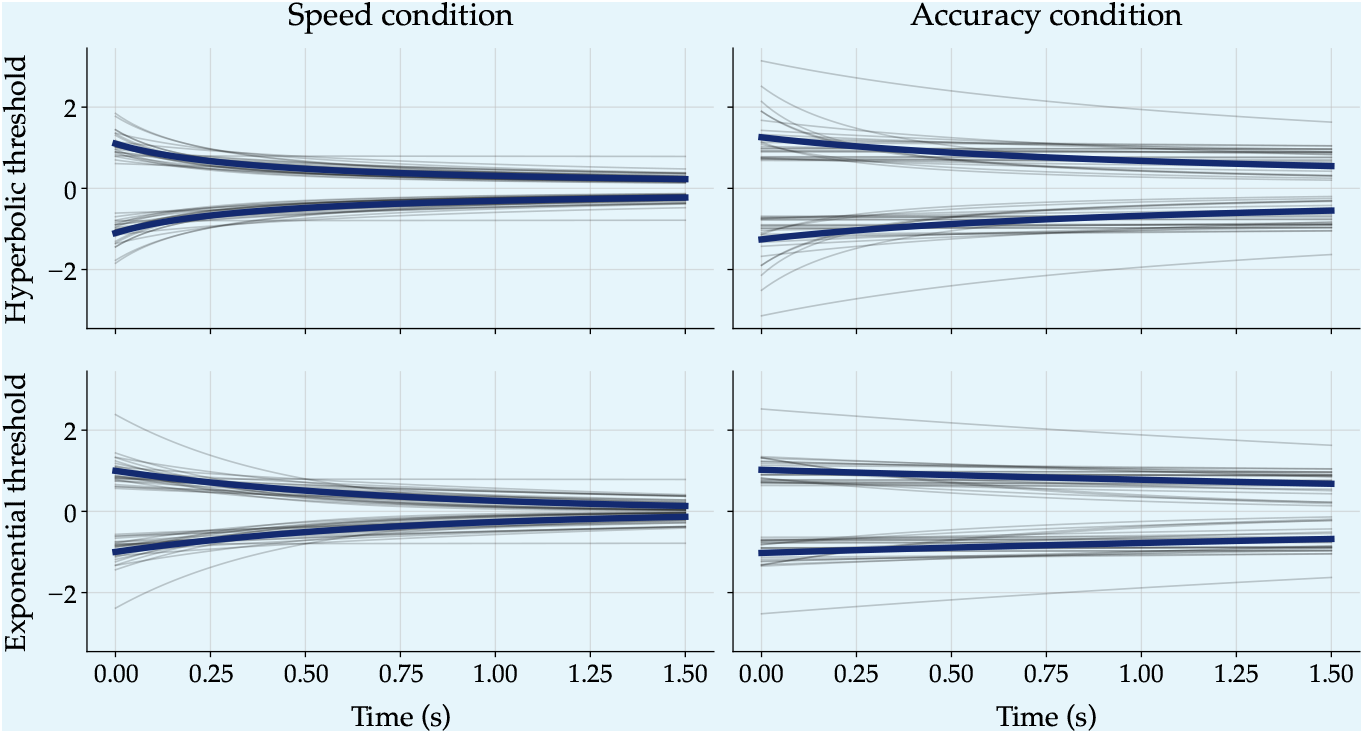
The estimated threshold dynamics from uninformed CT-DDMs for individuals in Study 1. The first row illustrates the estimated hyperbolic threshold dynamics, and the second row illustrates the estimated exponential threshold dynamics. The left column shows the threshold dynamics in the speed condition, and the right column shows the accuracy condition. Each gray line corresponds to a subject. The blue line corresponds to the average group level.

**Appendix 6—figure 2.**
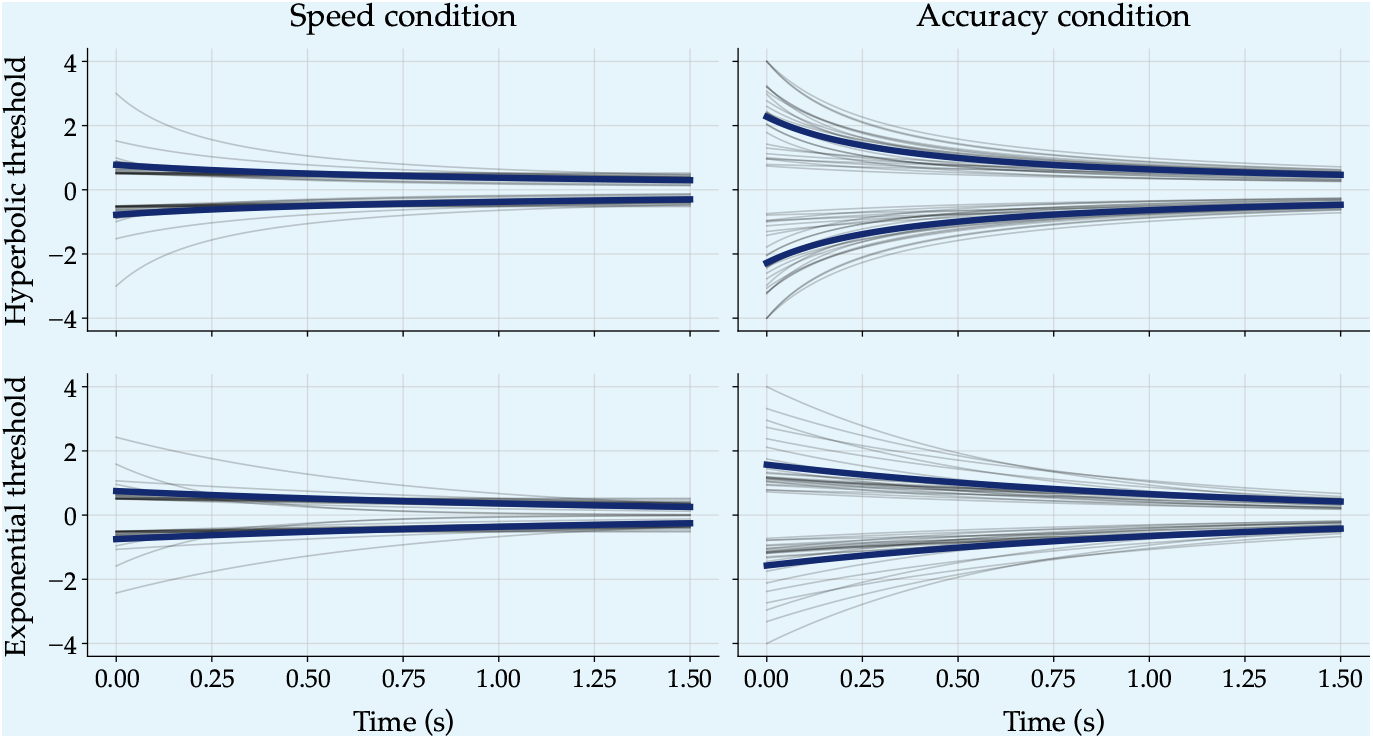
The estimated threshold dynamics from uninformed CT-DDMs for individuals in Study 2. The first row illustrates the estimated hyperbolic threshold dynamics, and the second row illustrates the estimated exponential threshold dynamics. The left column shows the threshold dynamics in the speed condition, and the right column shows the accuracy condition. Each gray line corresponds to a subject. The blue line corresponds to the average group level.

## Appendix 7

### Results of Fixed-threshold diffusion model with drift variability

#### Study 1

Figure 1 presents the predictions of the uninformed FT-DDM, which includes trial-to-trial variability in drift rate. This model overestimates RT quantiles in the speed condition, particularly for the later response time quantiles for both correct and incorrect responses. However, the predictions of the FT-DDM align more closely with observed empirical data in the accuracy condition. Fitting of this model on the speed condition resulted BIC_Behavior_ = 5817.66 and on the accuracy condition BIC_Behavior_ = 16577.20. These BIC values are much larger than those of all NDT-informed and uninformed CT-DDMs considered in the main text. This implies that all CT-DDMs perform better than the FT-DDM, which accounts for across-trial variability in drift rate.

**Appendix 7—figure 1.**
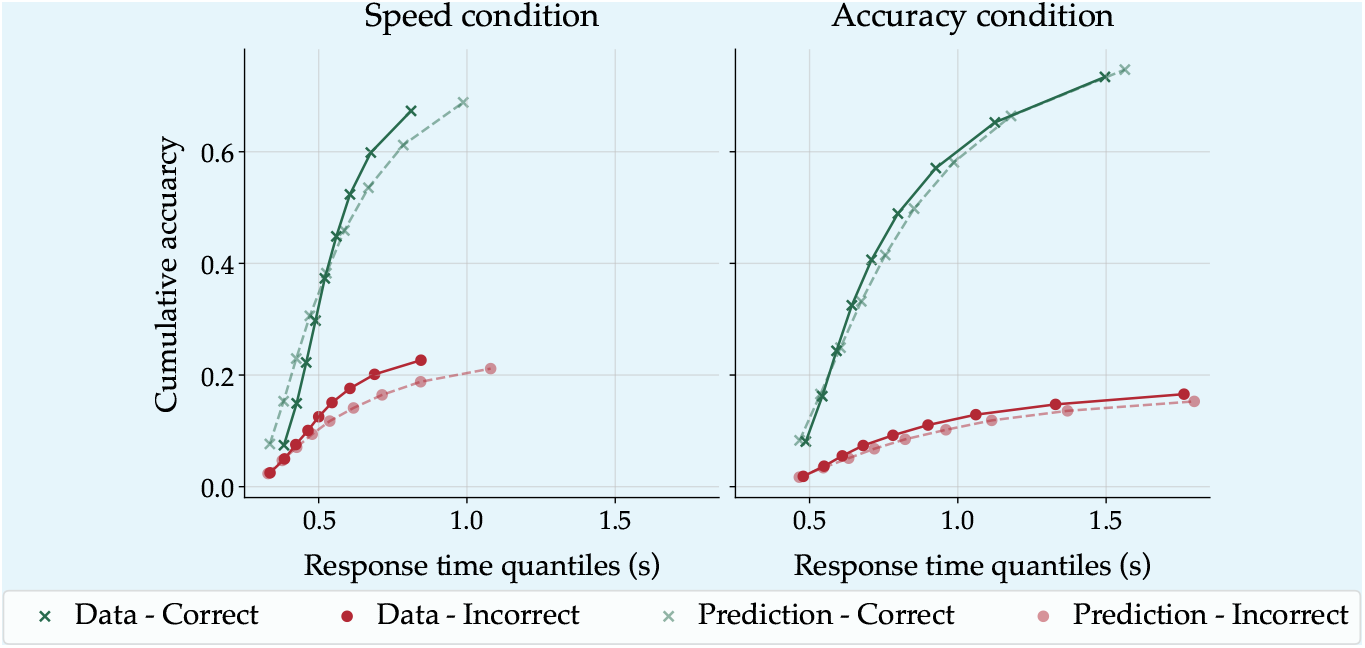
Posterior prediction of uninformed FT-DDM against empirical data for speed (left panel) and accuracy (right row) conditions in Study 1. In each panel, the x-axis represents the response time quantiles in seconds, and the y-axis represents the cumulative choice proportion.

#### Study 2

Similarly, the predictions of the FT-DDM with across-trial variability in drift rate are presented in Figure 2. The FT-DDM overestimates the RT quantiles for both correct and incorrect responses in both speed and accuracy conditions. Moreover, this model underestimates the proportion of incorrect responses chosen in the Speed condition. Also, the model fitting resulted in BIC_Behavior_ = 5855.32, which indicates that the fitting of CT-DDMs considered for study 2 in the main text is better than this model.

**Appendix 7—figure 2.**
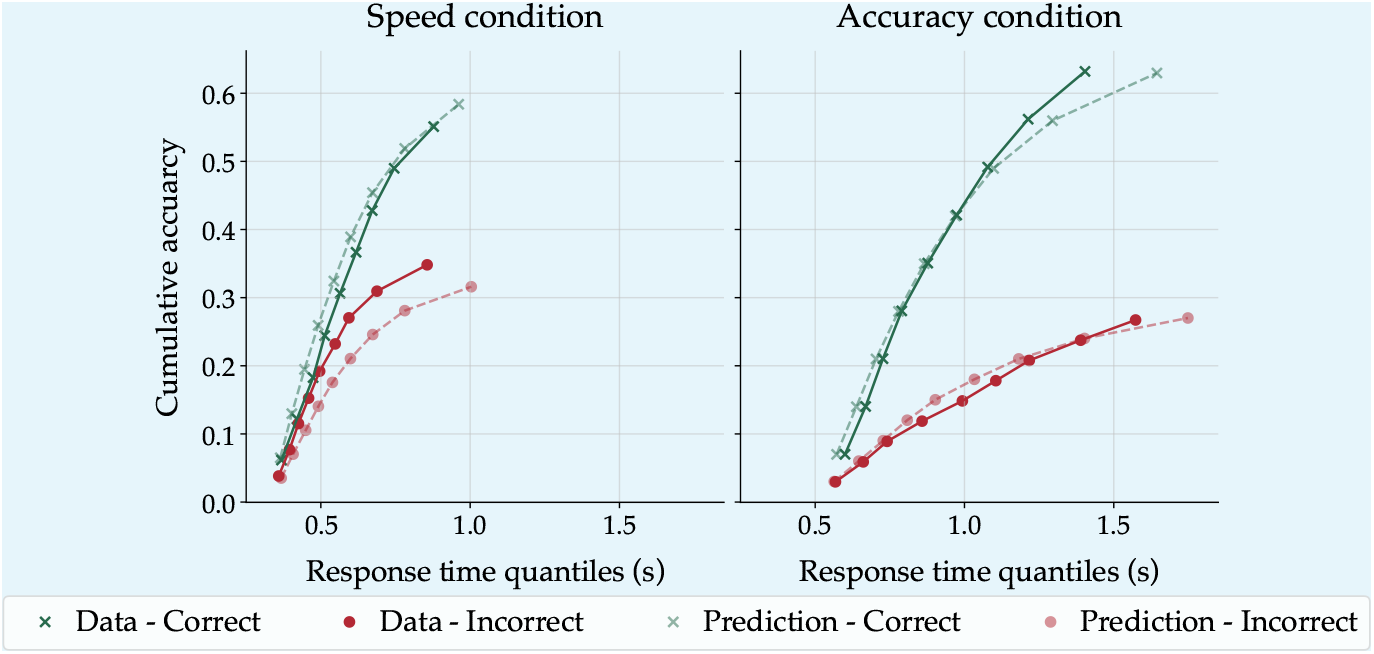
Posterior prediction of uninformed FT-DDM against empirical data for speed (left panel) and accuracy (right row) conditions in Study 2. In each panel, the x-axis represents the response time quantiles in seconds, and the y-axis represents the cumulative choice proportion.

## Appendix 8

### Simulation results for normally distributed non-decision time measurements

Non-decision time measurements can also be distributed normally around the actual non-decision time. This assumption has been made about the non-decision time variability in the literature (e.g., ***Ghaderi-Kangavari et al., 2023***; ***Kira et al., 2025***). Therefore, it is essential to demonstrate how the main results depend on the distributional assumptions underlying the non-decision time measurements. To this aim, we conducted a simulation study, in which we assumed that the response times and choices are generated from a CT-DDM, and the non-decision time measurements are normally distributed around the mean non-decision time *τ* with some variance 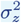 as follows:

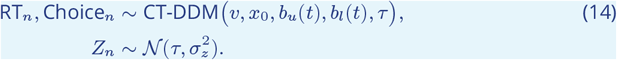

It is important to note that by considering the normal distribution for trial-level non-decision time measurements, the joint log-likelihood of response times, choices, and non-decision measurements becomes as follows:

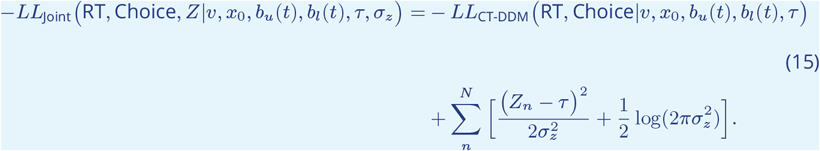

Figure 1 presents the *R*^2^ values for threshold parameters for NDT-informed models with normally distributed non-decision measurements against the uninformed models. Similar to the results reported in the main text with the log-normal assumption for non-decision time measurements, constraining the non-decision time parameter with normally distributed measurements also improves the recovery of the threshold parameters. Also, Figure 3 presents the estimated parameters against the actual generating parameters for both NDT-informed and uninformed models.

**Appendix 8—figure 1.**
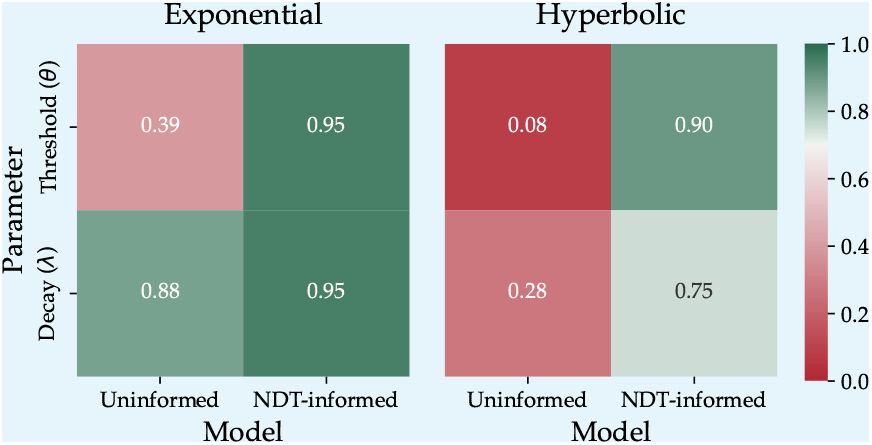
*R*^2^ values measuring the agreement between estimated and ground truth threshold parameters in exponential (left) and hyperbolic (right) collapsing boundary models.

**Appendix 8—figure 2.**
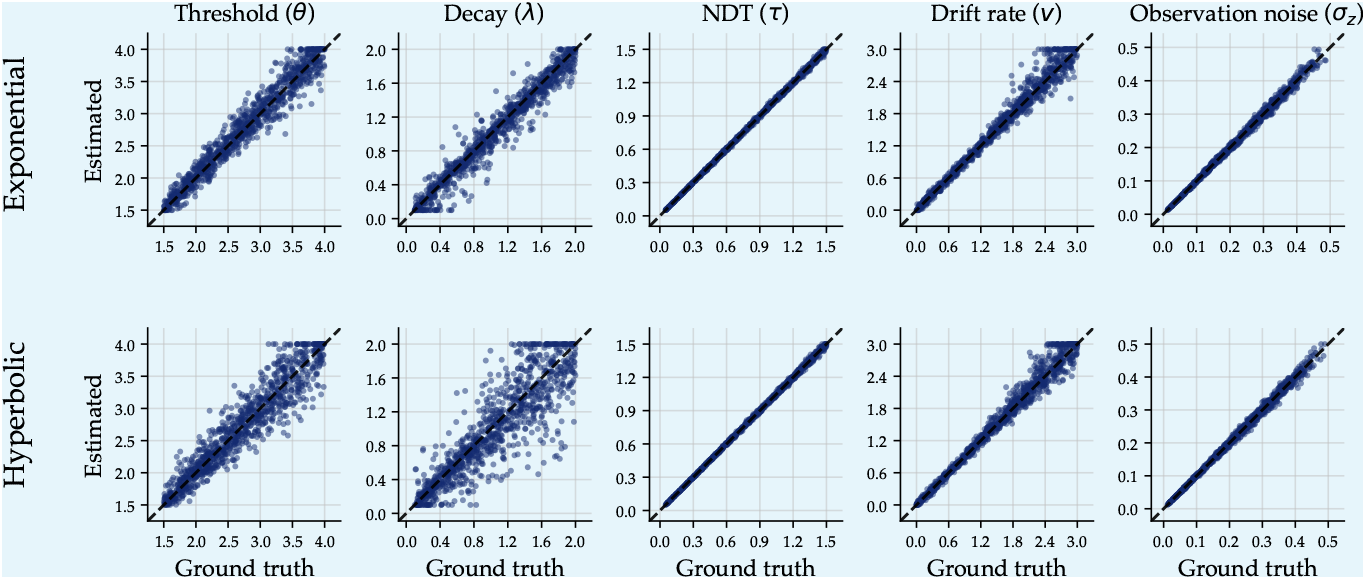
Estimated vs. true parameter values for two modeling approaches. The top row shows parameter recovery for the linear collapsing threshold diffusion model estimated without additional constraints (“Uninformed”), while the bottom row shows recovery for the joint model that incorporates additional non-decision time observations (“NDT informed”). Each point represents a recovered parameter estimate plotted against its corresponding true data-generating value. Dashed diagonal lines indicate perfect recovery.

Figure 3 illustrates the sensitivity of the parameter estimation to the number of trials. Similar to the results for the log-normal distribution, we also observed rapid improvement in parameter recovery with increasing the number of trials.

**Appendix 8—figure 3.**
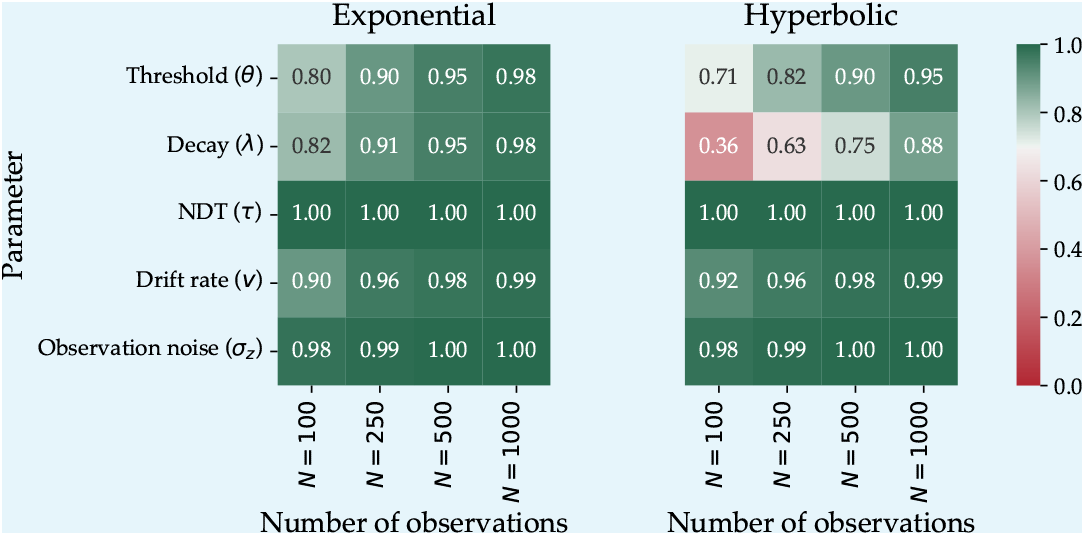
Illustration of the sensitivity of parameter estimation to the number of trials in the NDT-informed models. *R*^2^ values measure the agreement between estimated and ground truth parameters of the exponential (left) and hyperbolic (right) collapsing threshold models.

## Appendix 9

### Application to N-dimensional diffusion models

The hyper-spherical diffusion model (HSDM; ***Smith, 2016***; ***Smith and Corbett, 2019***; ***Hadian Rasanan et al., 2026***) is the *n*-dimensional extension of the well-known DDM (***Ratcliff, 1978***; ***Ratcliff and Rouder, 1998***). HSDM is an important model to consider because it is applicable in the modeling of decisions with continuous options (e.g., ***Smith et al., 2020, 2023***; ***Hadian Rasanan et al., 2026***), multi-alternative (e.g., ***Kvam, 2019a***; ***Smith and Corbett, 2019***), and multi-attribute decisions (e.g., ***Smith, 2019***; ***Smith and Corbett, 2019***). Thus, it covers a vast range of choice problems (see also ***Hadian Rasanan et al., 2024a***). Recently, ***Hadian Rasanan et al. (2025***) have extended the collapsing threshold version of HSDM and proposed an integral equation method for parameter estimation of this model. The evidence accumulation process in this model can be represented by an *n*-dimensional vector-valued Wiener process as follows:

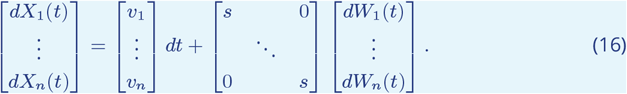

***Hadian Rasanan et al. (2025***) showed that the square distance of the an *n*-dimensional zero-drift process (i.e., *v*_*i*_ = 0 for *i* = 1, …, *n*) from the origin (i.e., 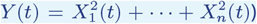 satisfies the following one-dimensional Feller-type stochastic differential equation (see also ***Göing-Jaeschke and Yor, 2003***; ***Kersting et al., 2023***):

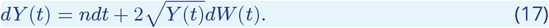

In this equation, *n* (number of dimensions) is the drift rate, and 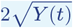 is the diffusion co-efficient. The authors showed that the first-passage time distribution of an *n*-dimensional zero-drift process starting from *y*_0_ in the presence of a time-dependent threshold *b*(*t*) satisfies the following integral equation (***Hadian Rasanan et al., 2025***):

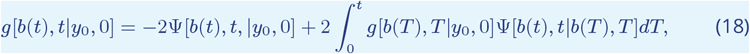

where the kernel function Ψ is defined as follows (***Giorno et al., 1989***; ***Hadian Rasanan et al., 2025***):

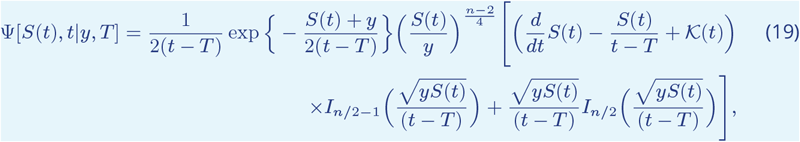

in which *S*(*t*) = *b*^2^(*t*), and 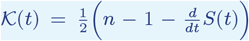. Also, *I*_*ς*_ (*z*) represents the Bessel function of the second kind of order *ς* defined as below (***Bell, 2004***):

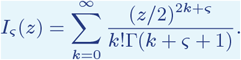

Similar to the one-dimensional CT-DDM, we need to approximate the solution of this integral equation to estimate the first-passage time distribution of the zero-drift process (see Appendix 2). For this purpose, we employed the following discretized scheme, which is identical to the approximation scheme employed for one-dimensional DDM (***Buonocore et al., 1987***; ***Hadian Rasanan et al., 2025***):

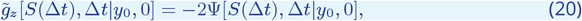

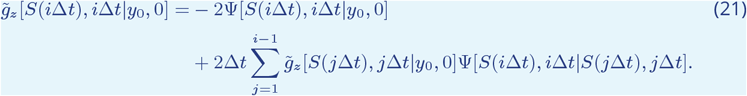

After estimating the first-passage time distribution of the zero-drift process, we apply the Girsanov change-of-measure theorem (***Girsanov, 1960***) to obtain the joint distribution of choice and RT for the non-zero drift process (***Smith, 2016***; ***Hadian Rasanan et al., 2025***). Therefore, the joint distribution of RT and choice can be obtained as follows:

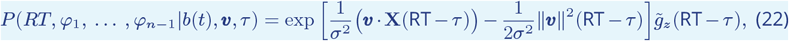

in which *v* = [*v*_1_, …, *v*_*n*_]^*′*^ is the drift vector, *φ*_*i*_ (*i* = 1, …, *n*− 1) are the response angles with respect to the *i*-th axis, and **X**(RT − *τ* ) denotes the final position of the accumulator in an *n-*dimensional space. The following Cartesian representation can be utilized for calculating the elements of the final stopping point **X**(RT − *τ* ):

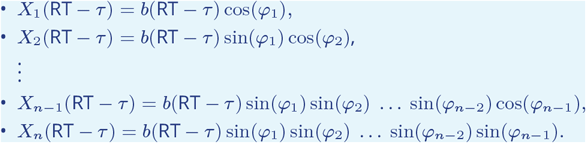

To assess the generalizability of the proposed NDT-informed diffusion modeling, we conducted a parameter recovery study based on the two-dimensional circular diffusion model (***Smith, 2016***; ***Hadian Rasanan et al., 2025***). Similar to the simulation study reported in the main text, we considered NDT-informed and uninformed collapsing threshold circular diffusion models (CT-CDM) with hyperbolic and exponential collapsing dynamics. We sampled 1000 random parameter sets from the following uniform distributions and then generated 500 trials from CT-CDM for each parameter set:

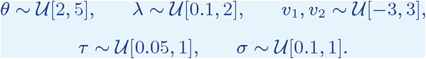

Figures 1 and 2 present the simulation results. Similar to the one-dimensional DDM, we also observe that constraining the non-decision time significantly improves parameter recovery of the two-dimensional CT-CDM.

**Appendix 9—figure 1.**
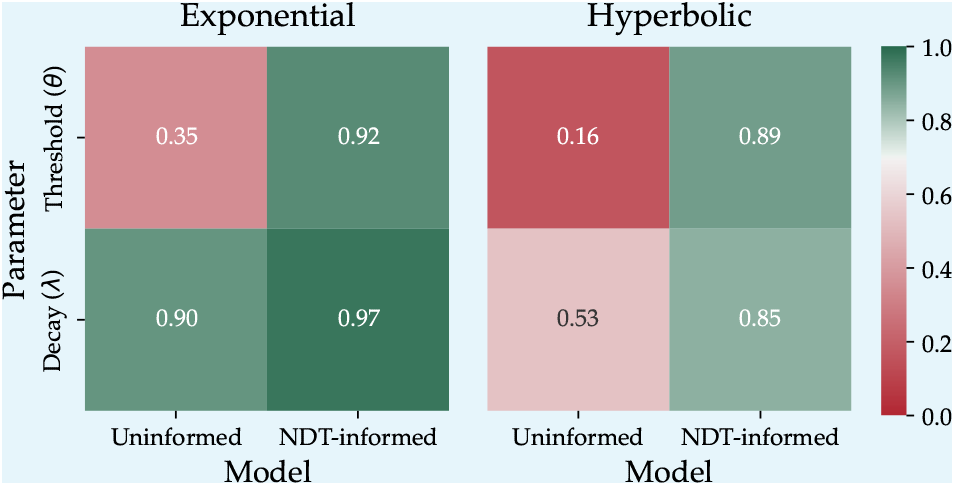
*R*^2^ values measuring the agreement between estimated and ground truth threshold parameters in exponential (left) and hyperbolic (right) collapsing boundary circular diffusion models.

**Appendix 9—figure 2.**
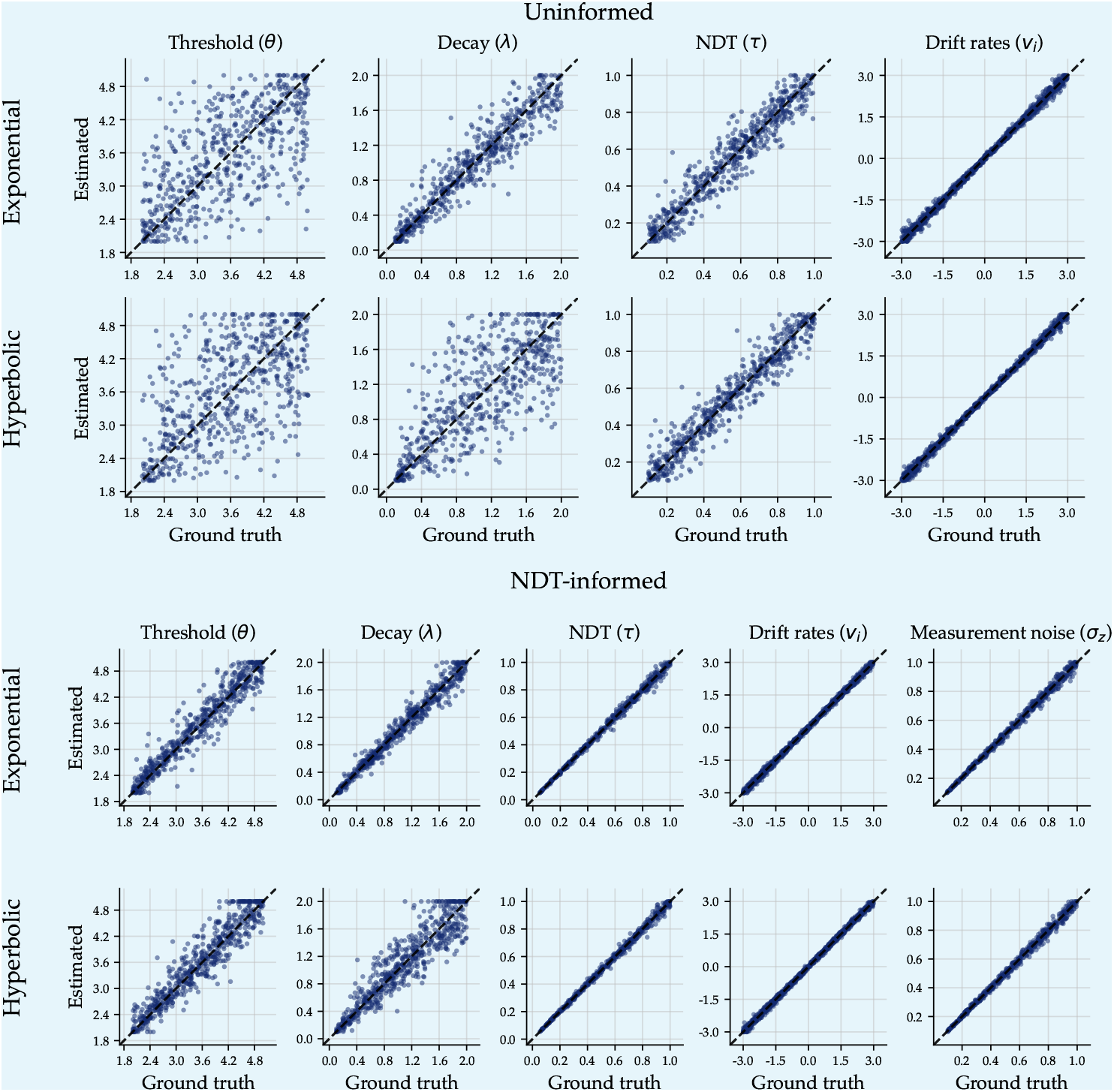
Estimated vs. true parameter values for two modeling approaches based on the CT-CDM. The top two rows show parameter recovery for models estimated without additional constraints (“Uninformed”), while the bottom two rows show recovery for joint models that incorporate additional non-decision time observations (“NDT-informed”). Each point represents a recovered parameter estimate plotted against its corresponding true data-generating value. Rows correspond to different functional forms (Exponential and Hyperbolic). Dashed diagonal lines indicate perfect recovery.

## Footnotes

1 Although the urgency gating model and collapsing threshold diffusion model are theoretically distinct, ***Smith and Ratcliff*** (***2022***) demonstrated that they can be transformed into one another. Therefore, we primarily focus on collapsing threshold models for the remainder of this paper; however, we will explain how the current work can also be related to urgency gating models in more detail in the general discussion.

2 It is worth noting that including trial-to-trial variability in drift rate enables the FT-DDM to predict slow errors (***Ratcliff and Rouder, 1998***; ***Voskuilen et al., 2016***).

3 Despite all the aforementioned evidence supporting CT-DDMs, some computational modeling studies have not found substantial improvements in model fit when incorporating a time-dependent threshold compared to a fixed threshold (e.g., ***Voskuilen et al., 2016***; ***Smith and Ratcliff, 2022***).

4 It is worth highlighting that this distributional assumption is about the non-decision time measurements and not about across-trial variability in non-decision time.

5 See also ***Kelly et al. (2021***) for a discussion on how to extract perceptual encoding onset, evidence accumulation onset, and post-decision motor-execution time from EEG signals.

6 Some studies have also employed the Weibull function for modeling threshold dynamics (e.g., ***Hawkins et al., 2015***; ***Ging-Jehli et al., 2025***; ***Evans et al., 2020a***). This choice is typically made for its flexibility, allowing the model to mimic a variety of threshold dynamical forms. However, this function has three parameters, which causes some trade-off among the threshold parameters and makes the parameter recovery much more difficult.

7 𝒰 [*a, b*] represent a uniform distribution over the interval between *a* and *b*.

8 As the number of parameters in both considered models is identical (i.e., the complexity of the two models is the same), we compared the log-likelihood value as a measure of goodness-of-fit to classify the underlying generative model.

9 Please note that, as the actual generating decay rate value for the FT-DDM is always zero, we employed a density plot to illustrate the density of the estimated decay rate.

10 However, we should also note that model recovery is highly sensitive to the experimental design considered. For instance, the experimental designs that include a condition with a high level of difficulty (i.e., the drift rate close to zero), then these models are more distinguishable.

11 The third aim is set due to the controversial findings in the literature that did not find evidence in support of CT-DDMs (e.g., ***Voskuilen et al., 2016***; ***Smith and Ratcliff, 2022***; ***Milosavljevic et al., 2010***; ***Karşılar et al., 2014***).

12 Since some studies showed the drift rate can be affected by speed/accuracy manipulation (e.g., ***Ratcliff and McKoon, 2023***), we also tested free-drift rate models in which the drift rate can vary across speed/accuracy conditions, and the obtained results were similar to what we have reported here.

